# GABA-independent activation of GABAB receptor by mechanical forces

**DOI:** 10.1101/2025.06.09.658551

**Authors:** Yujia Huo, Yiwei Zhou, Li Lin, Fan Yang, Jiyong Meng, Feiteng He, Fengfan Zhang, Mengdan Song, Cangsong Shen, Yuxuan Liu, Philippe Rondard, Chanjuan Xu, X.Z. Shawn Xu, Jianfeng Liu

## Abstract

The heterodimeric GABA_B_ receptor, composed of GB1 and GB2 subunits, is a metabotropic G protein-coupled receptor (GPCR) activated by the neurotransmitter GABA. GABA binds to the extracellular domain of GB1 to activate G proteins through GB2. Here we show that GABA_B_ receptors can be activated by mechanical forces, such as traction force and shear stress, in a GABA-independent manner. This GABA-independent mechano-activation of GABA_B_ receptor is mediated by a direct interaction between integrins and the extracellular domain of GB1, indicating that GABA_B_ receptor and integrin form a novel type of mechano-transduction complex. Mechanistically, shear stress promotes the binding of integrin to GB1 and induces an allosteric re-arrangement of GABA_B_ receptor transmembrane domains towards an active conformation, culminating in receptor activation. Furthermore, we demonstrate that shear stress-induced GABA_B_ receptor activation plays a crucial role in astrocyte remodeling. These findings reveal a previously unrecognized function of GABA_B_ receptor in mechano-transduction, uncovering a novel ligand-independent activation mechanism for GPCRs.

## Introduction

γ-Aminobutyric acid (GABA) is the primary inhibitory neurotransmitter in the brain. GABA_B_ receptor is a G protein coupled receptor (GPCR) that binds to GABA^1, 2^. As a metabotropic receptor, GABA_B_ receptor plays a crucial role in regulating brain functions and is implicated in various physiological processes, such as locomotion, learning and memory, and cognition^3–7^. It facilitates slow and long-term synaptic inhibition by inhibiting voltage-gated Ca^2+^ channels and preventing neurotransmitter release at presynaptic sites^8^. Additionally, it activates G protein-gated inwardly rectifying K^+^ channels and induces slow inhibitory potentials at postsynaptic sites in neurons^8^. GABA_B_ receptor also has the ability to trigger Ca^2+^ release^9, 10^ and regulate astrocyte morphogenesis^10, 11^. Dysregulation of GABA_B_ receptor activity has been implicated in various diseases, such as epilepsy^12, 13^, anxiety^14^, depression^15, 16^, spasticity^17^, addiction^18^, pain^19^, Rett-like Phenotype^20^, epileptic encephalopathy^21^ and fragile X syndrome^22^. Therefore, GABA_B_ receptor represents an excellent therapeutic target. Baclofen (Lioresal) is a specific agonist of GABA_B_ receptor and is used to treat spasticity in multiple sclerosis and alcohol abuse disorder^23–25^. Notably, GABA_B_ receptor is present not only in the central nervous system but also in peripheral regions such as the gastrointestinal (GI) tract, where it is involved in regulating intestinal motility, gastric emptying, and esophageal sphincter relaxation^26–28^. This raises the possibility that mechanical forces may regulate GABA_B_ receptor activity. However, whether mechanical forces can regulate GABA_B_ receptor activity has not been explored.

GABA_B_ receptor is a Class C GPCR that forms an obligatory heterodimer consisting of two subunits, GB1 and GB2. Each subunit comprises a large extracellular Venus flytrap domain (VFT) followed by canonical seven-transmembrane domains (7TMs)^29, 30^. The VFT domain consists of two lobes: LB1 and LB2^30^. In the inactive state, GABA_B_ receptor interacts through the LB1 of two subunits and the intracellular tips of transmembrane domain 3 and 5 (TM3 and TM5)^31^. Agonists (GABA, Baclofen) and antagonists (CGP54626) bind to a large crevice between LB1 and LB2 in the VFT domain of GB1 (VFT_GB1_). While the agonists stabilize the closure of VFT_GB1_, the antagonists prevent the closure of VFT_GB1_ (**Fig. 1a**). Agonist binding brings LB1 of VFT_GB1_ (LB1_GB1_) closer to LB2 of VFT_GB1_ (LB2_GB1_) to stabilize VFT_GB1_ closure, further bringing LB2_GB1_ and LB2 of GB2 (LB2_GB2_) in contact^30, 32–34^. The interplay between LB2_GB1_ and LB2_GB2_, which is essential for receptor activation, triggers a rearrangement from the inactive 7TM interface formed by intracellular tips of TM3 and TM5 of GB1 and GB2 to the active interface formed by TM6 of GB1 and TM6 of GB2, leading to the coupling of G_i/o_ proteins to GB2 in a shallow pocket formed by TM3 and three intracellular loops^30, 32–35^ (**Fig. 1a**). Additionally, positive allosteric modulators (PAMs) of GABA_B_ receptor, such as Rac BHFF, GS39783 and CGP7930, bind within the TM6-TM6 interface to stabilize the active conformation of the receptor^30, 32, 33, 36^. This mode of GABA_B_ receptor activation has been extensively characterized and validated using multiple approaches. However, it is unclear whether additional mechanisms are in place to activate GABA_B_ receptor.

**Fig. 1.**
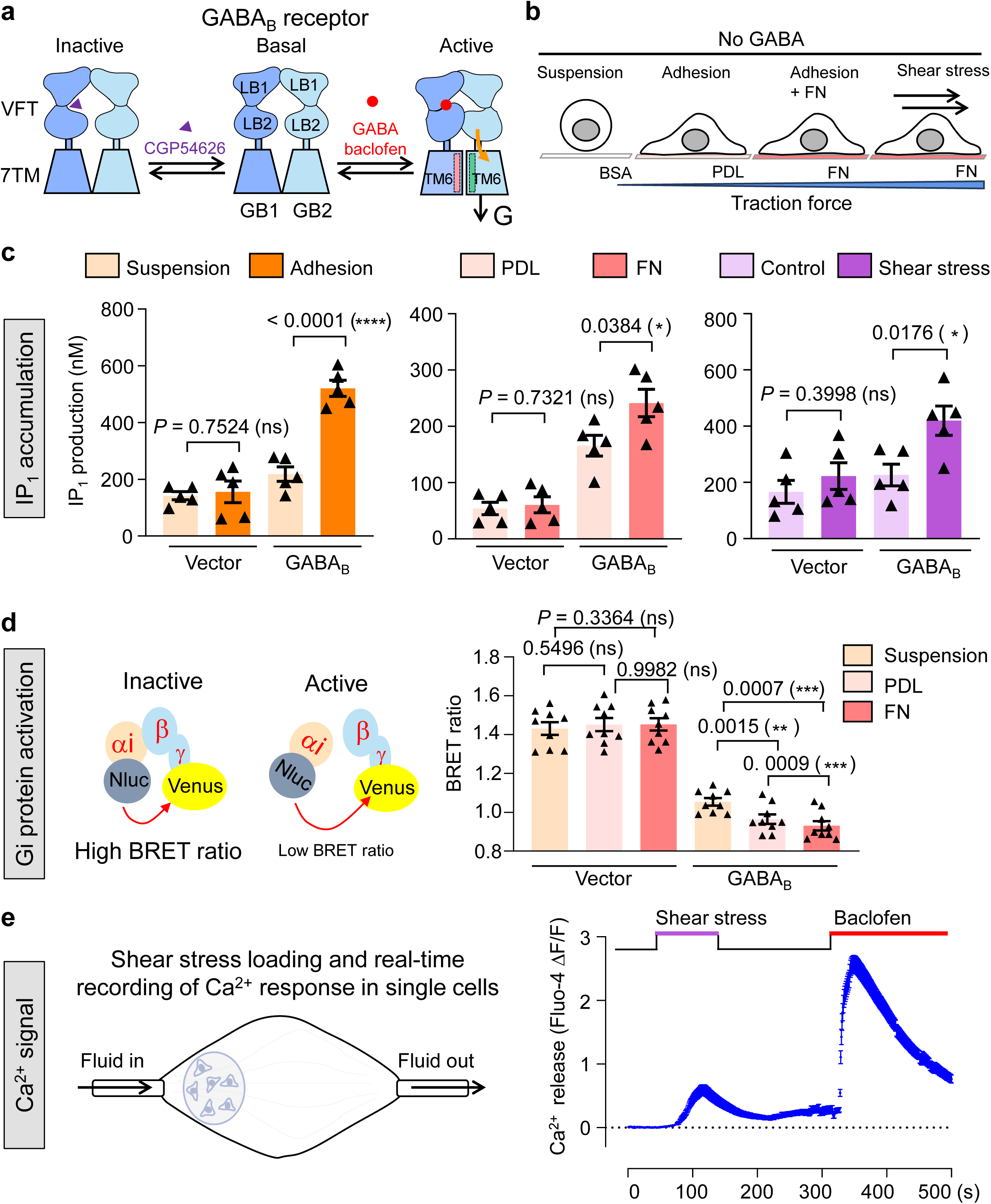
| Traction force and shear stress activate GABA_B_ receptor. **a**, Schematic illustration depicting the modulation of GABA_B_ receptor activity by ligands (e.g. agonists: GABA and baclofen; antagonist: CGP54626). **b,** Schematic representation of the experiments applying traction force and shear stress to cells in this study, encompassing conditions such as cell suspension or adhesion; PDL or FN coating; and shear stress treatment. No GABA was added in these experiments. **c,** IP_1_ production in HEK293 cells transfected with vector or GABA_B_ receptor, along with G_qi9_ chimera in the corresponding conditions: suspension or adhesion *(left)*; PDL or FN coating *(middle)*, without (control) or with shear stress (15 dyn/cm^2^, 15 min) *(right)*. Data are present as mean ± s.e.m. from five biologically independent experiments each performed in triplicates and analyzed using unpaired t test (two-tailed) to determine significance. \*\*\**P* < 0.001, \**P* < 0.05, not significant (ns) > 0.05. **d,** G_i_ protein activation in HEK293 cells transfected with vector or GABA_B_ receptor, along with G_i_ protein sensors under suspension, PDL-coating or FN-coating condition. Left: Schematic presentation of G_i_ protein activity measurement using G_i_ protein BRET sensor. Right: BRET ratio of cells in suspension or cells cultured in PDL-coating or FN-coating dishes. Data are present as mean ± s.e.m. from nine biologically independent experiments for vector and GABA_B_ receptor respectively each performed in triplicates, and analyzed using repeat measurements one-way ANOVA with Tukey’s multiple comparisons test to determine significance. \*\*\**P* < 0.001, \*\**P* < 0.01, not significant (ns) > 0.05. **e,** Real-time recording of intracellular Ca^2+^ release in HEK293 cells expressing GABA_B_ receptor and G_qi9_. Left: Schematic presentation of shear stress loading device and real-time recording of Ca^2+^ response in single cells. Right: Real-time Ca^2+^ signal measurement. After the recording of basal state of Ca^2+^ release for 50 seconds, cells were subjected to shear stress for 100 seconds. Shear stress was then halted for 150 seconds, after which baclofen was injected for 200 seconds. Data are present as mean ± s.e.m. from 85 cells recorded.

Here, we report a novel ligand-independent activation mechanism of the GABA_B_ receptor. We found that GABA_B_ receptor can be activated by mechanical forces (e.g., traction force and shear stress) independent of GABA. We demonstrated that GABA_B_ receptor and integrins form a novel type of mechano-transduction complex and that shear stress-induced GABA_B_ receptor activation plays a crucial role in astrocyte remodeling. Thus, our findings reveal a previously unrecognized function of the GABA_B_ receptor in mechano-transduction, uncovering a novel ligand-independent activation mechanism for GPCRs.

## Results

### GABA_B_ receptor is activated by mechanical forces

GABA_B_ receptor can be activated in a ligand-dependent manner by agonists, such as GABA and baclofen (**Fig. 1a**). However, whether a ligand-independent stimulus, such as mechanical forces, can activate GABA_B_ receptor remains unknown. To test this possibility, we explored whether different forces, such as traction force and shear stress, play a role in GABA_B_ receptor activation (**Fig. 1b**). Both traction force and shear stress are common mechanical forces experienced by cells^37, 38^. We first examined traction force. When growing on the surface of a substrate such as a culture dish coated with poly-D-lysine (PDL), cells usually adhere to the surface of the dish, and traction force develops between the cytoskeleton and extracellular matrix (ECM)^39^. However, no such traction force was observed in cells growing in suspension^40^. In addition, coating the culture dish with specialized ECM proteins, such as fibronectin (FN) which binds to the cell surface protein integrin, enhances traction force and promotes cell adhesion^41–43^. Myosin II activity is commonly used as a reliable readout of traction force in the cell^44, 45^. We found that in cultured HEK293 cells, myosin II activity was nearly undetectable in suspension cells; in contrast, myosin II activity was clearly observed in adherent cells growing on PDL-coated dishes (**Supplementary Fig. 1a**). Treatment with FN further increased myosin II activity in adherent HEK293 cells (**Supplementary Fig. 1b**). These results are consistent with the notion that traction force develops in adherent cells and further increases when cells were attached to ECM proteins^46^.

We then examined the activity of GABA_B_ receptor in both adherent and suspended HEK293 cells expressing GABA_B_ receptor and a chimeric Gq_i9_, in which the G_i/o_-coupled GABA_B_ receptor can couple to the PLC pathway^36^. We thereby measured the production of the downstream metabolite inositol monophosphate (IP_1_), using a commonly used assay that reports GPCR activity^47^. Interestingly, when cells expressing GABA_B_ receptor was attached on dishes coated with PDL, the IP_1_ production was significantly increased compared with that in suspension condition (**Fig. 1c**). The activity of GABA_B_ receptor was further increased in adherent cells growing on FN-coated dishes compared with PDL-coated dishes (**Fig. 1c**). Disruption of traction force by blebbistatin, which inhibits myosin II cross-bridge cycling^48^, blocked GABA_B_ receptor activity in adherent cells (**Supplementary Fig. 1c**). These results reveal a correlation between the GABA_B_ receptor activity and traction force, suggesting that mechanical forces can activate GABA_B_ receptor. Because FN coating reflects a more physiological condition, we decided to focus on characterizing cells growing under this condition. Next, we evaluated the effect of shear stress on GABA_B_ receptor. Shear stress increased IP_1_ production in adherent HEK293 cells expressing the GABA_B_ receptor and Gq_i9_ but not in control cells expressing only Gq_i9_ (**Fig. 1c**). This experiment suggests that shear stress can also activate GABA_B_ receptor.

We also performed experiments to verify if GABA_B_ receptor-induced direct G_i/o_ protein responses can be activated by traction force, using a sensitive BRET G_i_ protein sensor as reported by us recently^49^. The decreased BRET ratio represents the conformational rearrangement between Gα_i_ and Gβγ subunit and the activation of G_i_ protein. The BRET ratio in GABA_B_ receptor-expressing cells was lower than that in control cells expressing only G_i_ protein sensor (**Fig. 1d**), due to the constitutive activity of GABA_B_ receptor^20, 33, 36^. The BRET signal was significantly decreased in cells expressing GABA_B_ receptor cultured on PDL or FN-coated surface compared with that in suspension condition (**Fig. 1d**). As a control, the BRET ratio in cells expressing only G_i_ protein sensor (**Fig. 1d**), or expressing a GABA_B_ receptor mutant which cannot couple to G protein (GABA_B_-ΔG) (**Supplementary Fig. 1d**), show no significant change compared with suspension, PDL-coating and FN-coating condition. This experiment confirms that mechano-force activates GABA_B_ receptor-induced G_i/o_ signaling.

While IP_1_ production serves as a reliable readout for GABA_B_ receptor activation, it only reports the accumulative rather than the dynamic activity of GABA_B_ receptor. To provide further evidence, we performed calcium imaging experiments to monitor the dynamic activity of GABA_B_ receptors in response to shear stress in real-time. We found that shear stress could also induce GABA_B_ receptor activity in cells expressing GABA_B_ receptor and G_qi9_ shown by the calcium imaging assay (**Fig. 1e**). A transient increase of Ca^2+^ release was observed after shear stress application, which quickly peaked and then gradually decreased to basal levels (**Fig. 1e**). The kinetics of shear stress-induced Ca^2+^ were slower, and the strength of shear stress-induced Ca^2+^ was lower than that of baclofen (**Supplementary Fig. 1e**). No such activity was observed in the control cells expressing only G_qi9_ (**Supplementary Fig. 1f**). Thus, both assays reveal that shear stress can activate GABA_B_ receptor. Taken together, these results demonstrated that GABA_B_ receptor can be activated by mechanical forces. No GABA or any other GABA_B_ receptor agonist was present in our assays and the concentration of GABA in the medium or the buffer was very low (**Supplementary Fig. 2**), indicating that the observed mechano-activation of the GABA_B_ receptor is ligand-independent.

### Mechano-activation of GABA_B_ receptor occurs through a physical interaction with integrin

How is GABA_B_ receptor activated by mechanical forces? The observation that the GABA_B_ receptor can be activated in adherent cells but not in suspension cells indicates that the mechano-activation of GABA_B_ receptor is cell condition-specific, suggesting the participation of additional proteins. As the ECM protein FN greatly enhanced the mechano-activation of GABA_B_ receptor (**Fig. 1c**) and FN acts by directly binding to the cell surface protein integrin^50^, we hypothesized that integrin may be involved in the mechano-activation of GABA_B_ receptor. The fact that integrin is a mechano-sensor, which connects both ECM proteins and the cytoskeleton and is sensitive to traction force and shear stress^51, 52^, further supports this model. To test this model, we turned our attention to integrin β_3_, which is the primary receptor for FN^53, 54^. As a receptor of FN, integrin β_3_ can function as a heterodimer with integrin α ^55^. Knockdown of integrin β_3_ by siRNA abolished traction force-induced GABA receptor activation (**Fig. 2a, Supplementary Fig. 3a**). These results suggest that integrin β_3_ plays an important role in the mechano-activation of GABA_B_ receptor.

**Fig. 2.**
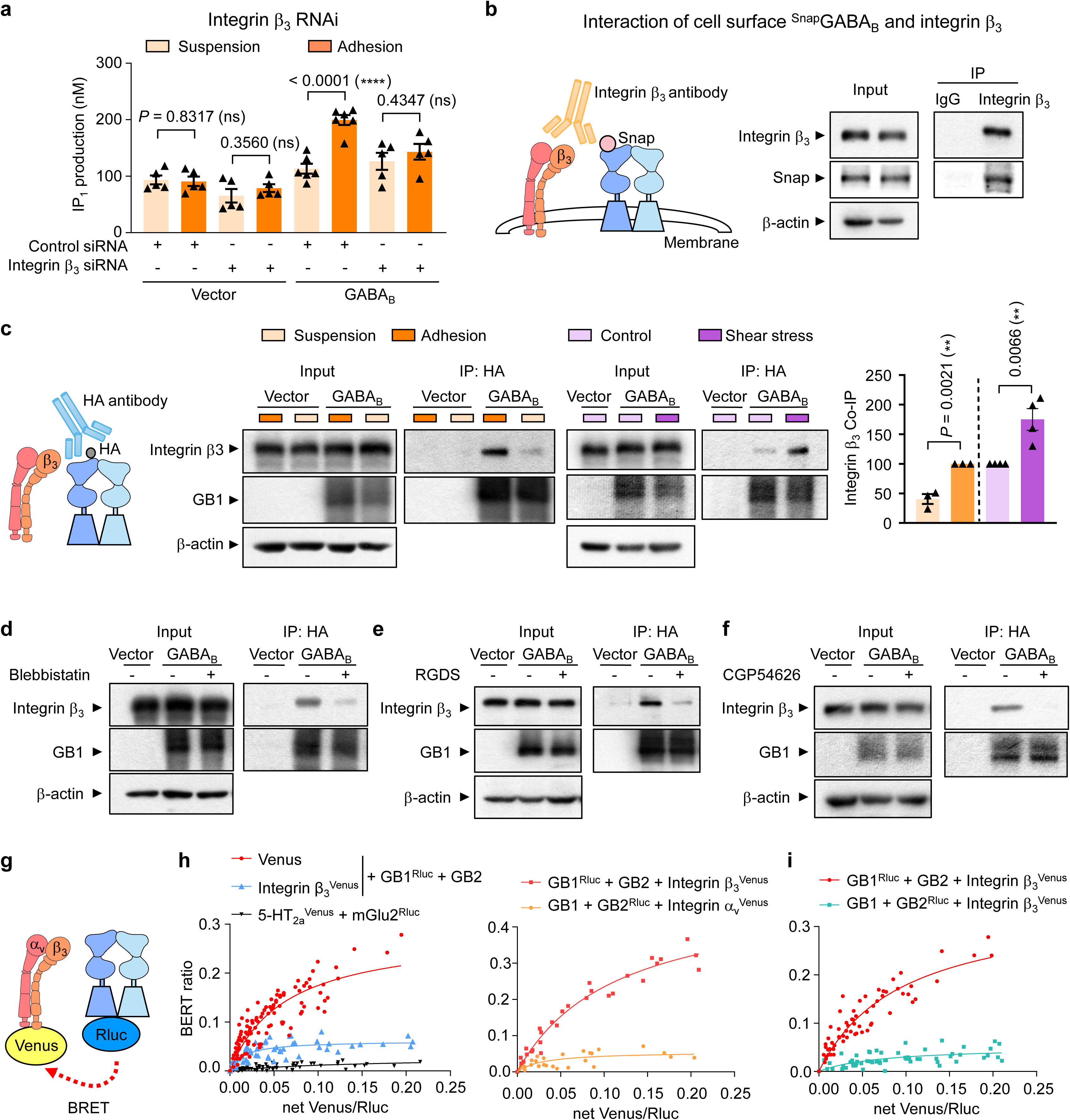
| Mechano-activation of GABA_B_ receptor requires integrin β_3_ interaction. **a**, IP_1_ production in HEK293 cells transfected with either control siRNA or integrin β3 siRNA, along with the expressing of vector and G_qi9_, or GABA_B_ receptor and G_qi9_, under either adhesion or suspension conditions. Data are present as mean ± s.e.m. from five biologically independent experiments and analyzed using unpaired t test (two-tailed) to determine significance. \*\*\**P* < 0.001, not significant (ns) > 0.05. **b**, Co-immunoprecipitation of GABA_B_ receptor and integrin β_3_ in HEK293 cells transfected with GABA_B_ receptor constructs (Snap-tagged GB1 and GB2) using anti-integrin β_3_ antibody, under basal condition. Only GB1 in cell surface was labeled and visualized using an impermeable Snap fluorescent substrate. **c-f,** Co-immunoprecipitation of GABA_B_ receptor and integrin β_3_ in HEK293 cells transfected with GABA_B_ receptor constructs (HA-tagged GB1 and GB2) using anti-HA antibody, under conditions including suspension or adhesion *(c)*, without (control) or with shear stress *(c)*, Blebbistatin (50 μM, 30 min) treatment *(d)*, RGDS (10 μM, 12 h) treatment *(e)*, or CGP54626 (50 μM, 30 min) treatment *(f)*. Blots are representative from at least three biologically independent experiments (*c*, suspension or adhesion, n = 3; control or with shear stress, n = 4; *d*, n = 3; *e*, n = 3; *f*, n = 4). The amount of integrin β_3_ immunoprecipitated by IgG or HA antibody are present as mean ± s.e.m. in *(e)* and analyzed using unpaired t test (two-tailed) to determine significance. \*\**P* < 0.01. **g,** Schematic representation of the BRET assay detecting direct interaction between GABA_B_ receptor and integrin β_3_. Rluc was fused in C-terminal of GB1 or GB2 subunit (GB1^Rluc^ or GB2^Rluc^) as luminescence donor. Venus was fused in C-terminal of integrin β_3_ or integrin α_V_ subunit (integrin β_3Venus_ or integrin α ^Venus^) as fluorescence acceptor. **h-i,** Interaction of GABA_B_ receptor and integrin β_3_ interaction between GB1 and integrin β_3_ or GB1 and integrin α_V_ *(h)*, or GB2 and integrin β_3_ *(i)* detected using BRET titration assay. Data were analyzed by nonlinear regression on a pooled data set from at least three biologically independent experiments each performed in triplicates, fitting with 1-site binding model in GraphPad Prism 8.

To investigate how integrin β_3_ mediates the mechano-activation of GABA_B_ receptor, we asked whether GABA_B_ receptor physically interacts with integrin β_3_. Indeed, when expressed in HEK293 cells, GABA_B_ receptor co-immunoprecipitated (co-IP) with endogenous integrin β_3_. This was performed using two different approaches to tag (HA tag and Snap tag) the GB1 subunit of GABA_B_ receptor (**Fig. 2b, Supplementary Fig. 3b-d**). We also detected integrin α_v_ in the complex **(Supplementary Fig. 3e)**, which is consistent with the notion that integrin β_3_ functions as a heterodimer with integrin α_v50, 56_. Thus, GABA_B_ receptor and integrin α_v_β_3_ appear to form a protein complex. In support of this, we found that GABA_B_ receptor and integrin β_3_ also co-localized in HEK293 cells (**Supplementary Fig. 3f**). These results together demonstrate GABA_B_ receptor physically interacts with integrin α_v_β_3_ to form a protein complex.

Interestingly, the interaction between integrin β_3_ and GABA_B_ receptor appeared to be much stronger in adherent cells **(Fig. 2c)**. Disruption of traction force using blebbistatin greatly diminished the interaction between integrin β_3_ and GABA_B_ receptor **(Fig. 2d, Supplementary Fig. 3g)**. Conversely, applying shear stress to HEK293 cells further enhanced integrin β_3_ and GABA_B_ receptor interaction in these cells (**Fig. 2c**,). These findings indicate that mechanical forces promote the formation of a GABA_B_ receptor-integrin α_v_β_3_ complex by facilitating the interaction between the two proteins.

To provide further evidence that integrin α_v_β_3_ and GABA_B_ receptor physically interact with each other, we tested the inhibitor and the antagonist. The integrin β_3_ inhibitor RGDS and the GABA_B_ receptor antagonist CGP54626 both inhibited the interaction between integrin β_3_ and GABA_B_ receptor (**Fig. 2e-f, Supplementary Fig. 3g**). This suggests that the interaction between integrin β_3_ and GABA_B_ receptor is rather specific. Furthermore, the formation of GABA_B_ receptor and integrin α_v_ β_3_ complex increased with GABA treatment (**Supplementary Fig. 3h**). Overall, our observation suggests that the formation of GABA_B_ receptor-integrin α_v_β_3_ complex may depend on their active conformation.

We then investigated whether the interaction between GABA_B_ receptor and integrin α_v_β_3_ is direct. To test this, we performed a bioluminescence resonance energy transfer (BRET) titration experiment^57, 58^. The energy donor Rellina luciferase (Rluc) and the energy acceptor Venus were attached to the C-terminus of GB1 or GB2 subunit of GABA_B_ receptor, integrin β_1_, integrin β_3_ and integrin α_v_ (named as GB1_Rluc_, GB2_Rluc_, integrin β_1Venus_, integrin β_3Venus_ and integrin α_vVenus_), respectively (**Fig. 2g**). Within a distance of 10 Å, energy transfer occurs between Rluc and Venus, which would indicate a direct interaction between the two proteins. In this assay, by examining the BRET ratio as a function of the relative amounts of the two tested proteins, one would not only be able to evaluate if a direct interaction occurs between the two proteins, but also can assess the relative strength of the interaction. We first performed control experiments. While no BRET signals were detected between Venus and GABA_B_ receptor (negative control), strong BRET signals were observed between mGlu2 and 5-HT_2a_ that are known to directly interact (positive control)^59^, validating the assay. Importantly, we observed strong BRET signals between GABA_B_ receptor and integrin β_3_ (**Fig. 2h**). These BRET signals were much more robust than those between mGlu2 and 5-HT_2a_, suggesting that the interaction between GABA_B_ receptor and integrin β_3_ is stronger than that between mGlu2 and 5-HT_2a_, which has been demonstrated to form oligomer^59^. By contrast, weak, if any, BRET signals were detected between integrin α_v_ and GABA_B_ receptor **(Fig. 2h)**. Thus, although integrin α_v_ was present in the integrin α_v_β_3_-GABA_B_ receptor complex **(Supplementary Fig. 3e)**, it did not appear to directly interact with GABA_B_ receptor, further supporting the model that integrin β_3_ rather than α_v_ directly interacts with GABA_B_ receptor. Notably, the BRET signals between GB1 and integrin β_3_ were much more robust than those between GB2 and integrin β_3_, suggesting that GABA_B_ receptor interacts with integrin β_3_ primarily via its GB1 subunit (**Fig. 2i**). These results demonstrate that the mechano-activation of GABA_B_ receptor requires integrin α_v_β_3_, which is primarily mediated by a direct interaction between integrin β_3_ and the GB1 subunit of GABA_B_ receptor. Taken together, our data suggest that GABA_B_ receptor responds to mechanical forces through a novel mechano-transduction complex formed with integrin α_v_β_3_ and that mechanical forces can promote the formation of this protein complex. FN can also bind to integrin α_v_β_160_. BRET signals were also observed between integrin β1 and GABA_B_ receptor (**Supplementary Fig. 3i**), suggesting a formation of mechano-transduction GABA_B_ receptor complex with FN-related integrin proteins. Interestingly, other ECM proteins such as collagen I also increased GABA_B_ receptor activation (**Supplementary Fig. 3j**), suggesting the involvement of other integrin subtypes.

### Both GB1 and GB2 are required for GABA_B_ receptor activation by mechanical forces

As GABA_B_ receptor interacted with integrin β_3_ primarily through its GB1 subunit, we wondered if GB1 alone can interact with integrin β_3_. As GB1 alone cannot localize to the plasma membrane, we tested GB1_ASA_, a GB1 subunit mutant which can localize to the cell surface on its alone^61^, and found that it cannot be activated by traction force (**Fig. 3a**). A similar phenomenon was observed with GB2 alone (**Fig. 3a**). Thus, both GB1 and GB2 subunits are required for the mechano-activation of GABA_B_ receptor. As GB2 is responsible for coupling G proteins to GABA_B_ receptor upon agonist binding, we tested GB2_L685P_, a mutant form of GB2 that is incapable of activating G proteins^62^. The traction force failed to induce GABA_B_ receptor activation in HEK293 cells expressing GB1, GB2_L685P_ and G_qi9_ (**Fig. 3b**). Thus, similar to ligand-dependent activation of GABA_B_ receptor, mechanical activation of this receptor also requires coupling of G proteins to the receptor via the GB2 subunit. These results indicate that while GABA_B_ receptor interacts with integrin β_3_ primarily via the GB1 subunit, the GB2 subunit is also necessary for GABA_B_ receptor activation by mechanical forces and this ligand-independent activation mode required G protein coupling to GB2. Thus, both GB1 and GB2 subunits participate in the formation of the mechano-transduction complex with integrin α_v_β_3_.

**Fig. 3.**
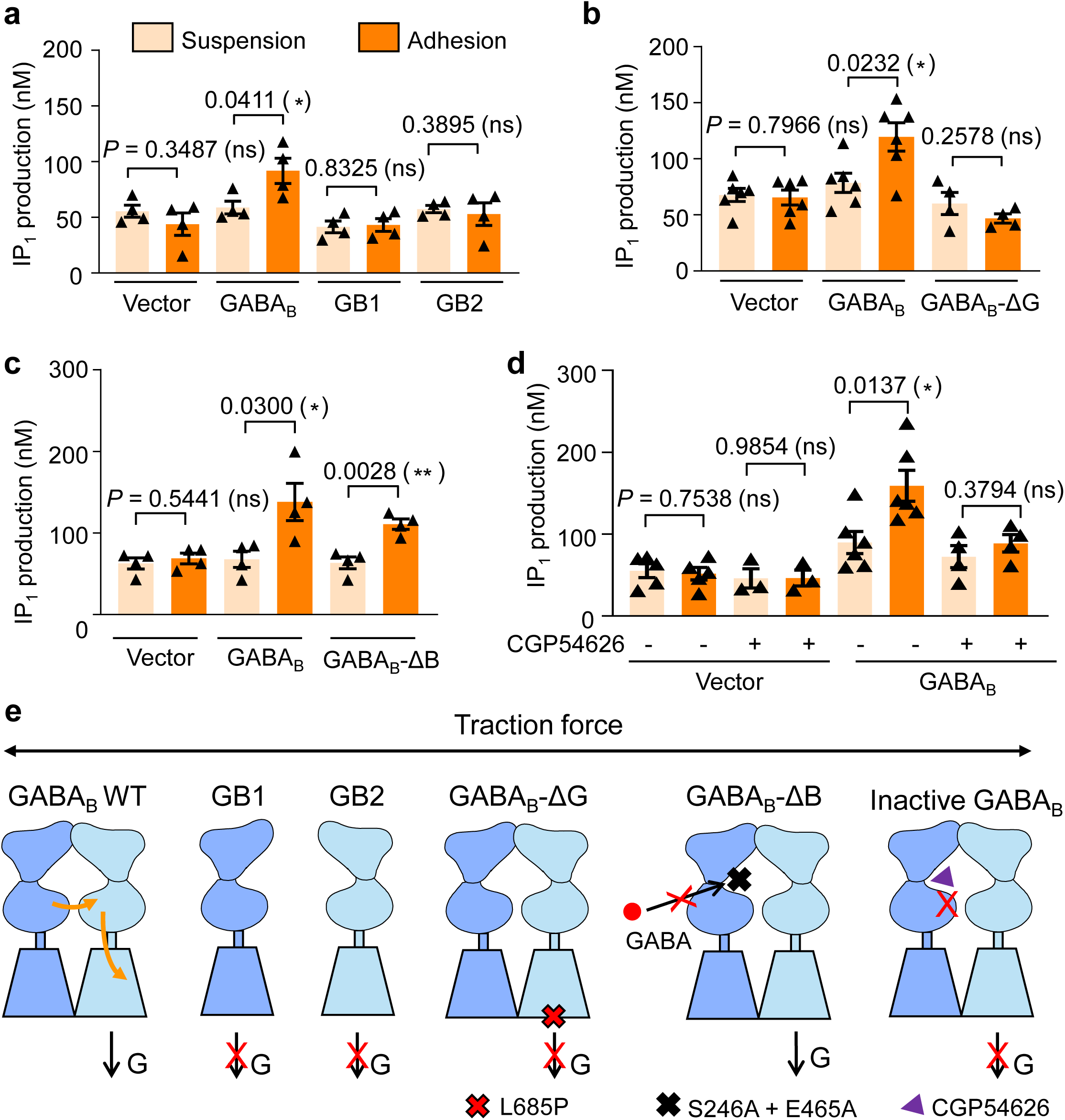
| Both GB1 and GB2 subunits are required for GABA_B_ receptor activation by mechanical forces. **a-d**, IP_1_ production in HEK293 cells transfected with the indicated constructs, along with G_qi9_ under either suspension or adhesion conditions: *(a)*: vector, GABA_B_ receptor (GB1 and GB2), GB1 only or GB2 only; *(b)*: vector, GABA_B_ receptor or GABA_B_-ΔG (mutant with L685 in GB2 substituted with Phenylalanine that fails to couple G protein); *(c)*: vector, GABA_B_ receptor, or GABA_B_-ΔB (mutant with the residues S276 and E465 in GB1 mutated to Analine that fails to bind GABA); *(d)*: vector or GABA_B_ receptor, treated with or without antagonist CGP54626 (50 μM, 30 min). Data are present as mean ± s.e.m. from at least three biologically independent experiments (n = 4, 6, 5, 5 for *a-d* respectively) each performed in triplicates, and analyzed using unpaired t test (two-tailed) to determine significance. \*\*\**P* < 0.001, \**P* < 0.05, not significant (ns) > 0.05. **e,** Model of the traction force-induced GABA_B_ receptor activation through the closure of GB1_VFT_-induced LB2_GB1_ and LB2_GB2_ in contact. The traction force-activated GABA_B_ receptor requires both GB1 and GB2, and relies on GB2 for G protein coupling. Whereas the traction force-activated GABA_B_ receptor is independent of GABA binding, preventing VFT_GB1_ closure by antagonist CGP54626 GABA_B_ receptor abolishs traction force-induced GABA_B_ receptor activation, indicating the closure of GB1_VFT_-induced LB2_GB1_ and LB2_GB2_ in contact is important for traction force-induced GABA_B_ receptor activation.

Furthermore, to understand how mechano-activation occurs through the interaction between the GB1 and GB2 subunits, we generated a GB1 mutant by inserting two mutations (S246A and E465A) into GB1_VFT_ as previously reported^63–65^ to prevent GABA binding but not to prevent the closure of GB1_VFT_-brought GB1_LB2_ and GB2_LB2_ in contact (**Fig. 3c**). The GABA_B_-ΔB composed by this GB1 mutant and GB2 wild type can still be activated by traction force to the similar extend as GABA_B_ receptor (**Fig. 3c**), demonstrating GABA-binding into GB1_VFT_ was not required in mechano-activation of GABA_B_ receptor and indicating that mechano-force alone also stabilize the closure of GB1_VFT_-brought LB2 of two subunits in contact. An antagonist (CGP54626) can prevent GB1_VFT_ closure-induced two LB2 in contact^32^. CGP54626 completely blocked traction force-induced GABA_B_ receptor activation, indicating that mechano-force induced GABA_B_ receptor activation requires the closure of GB1_VFT_-brought two LB2 of two subunits in contact (**Fig. 3d**). This data is also consistent with the observation that CGP54626 inhibited the interaction between integrin β_3_ and GABA_B_ receptor (**Fig. 2f)** whereas GABA_B_-ΔB still interacts with integrin β_3_ (**Supplementary Fig. 3k**). Overall, our results show that the traction force-activated GABA_B_ receptor requires both GB1 and GB2 subunit, and is dependent on the closure of GB1_VFT_-induced LB2_GB1_ and LB2_GB2_ in contact, and GB2 subunit for G protein coupling, but independent of GABA binding, suggesting that mechano-force acts as a positive allosteric modulator agonist (PAM Ago) of the GABA_B_ receptor (**Fig. 3e**).

### Mapping the region in GABA_B_ receptor that interacts with integrin β_3_

We then sought to map the region in GB1 that interacts with integrin β_3_. First, we first examined the role of VFT in the extracellular domain of GB1 (VFT_GB1_). Deleting VFT in GB1with a truncated GABA_B_ receptor (GABA_B_-ΔVFT) lacking the VFT domain of GB1 (GB1-ΔVFT) (**Fig. 4a**), abolished the interaction between integrin β_3_ and GABA_B_ receptor, revealing a key role of VFT_GB1_ in mediating the interaction **(Fig. 4b, Supplementary Fig. 4a**). Similarly, traction force and shear stress failed to induce GABA_B_-ΔVFT activation in HEK293 cells (**Fig. 4c-d**), indicating that the VFT_GB1_ is required for the mechano-activation of GABA_B_ receptor. Both results suggest that the VFT_GB1_ may mediate the physical interaction between GB1 and integrin β_3_.

**Fig. 4.**
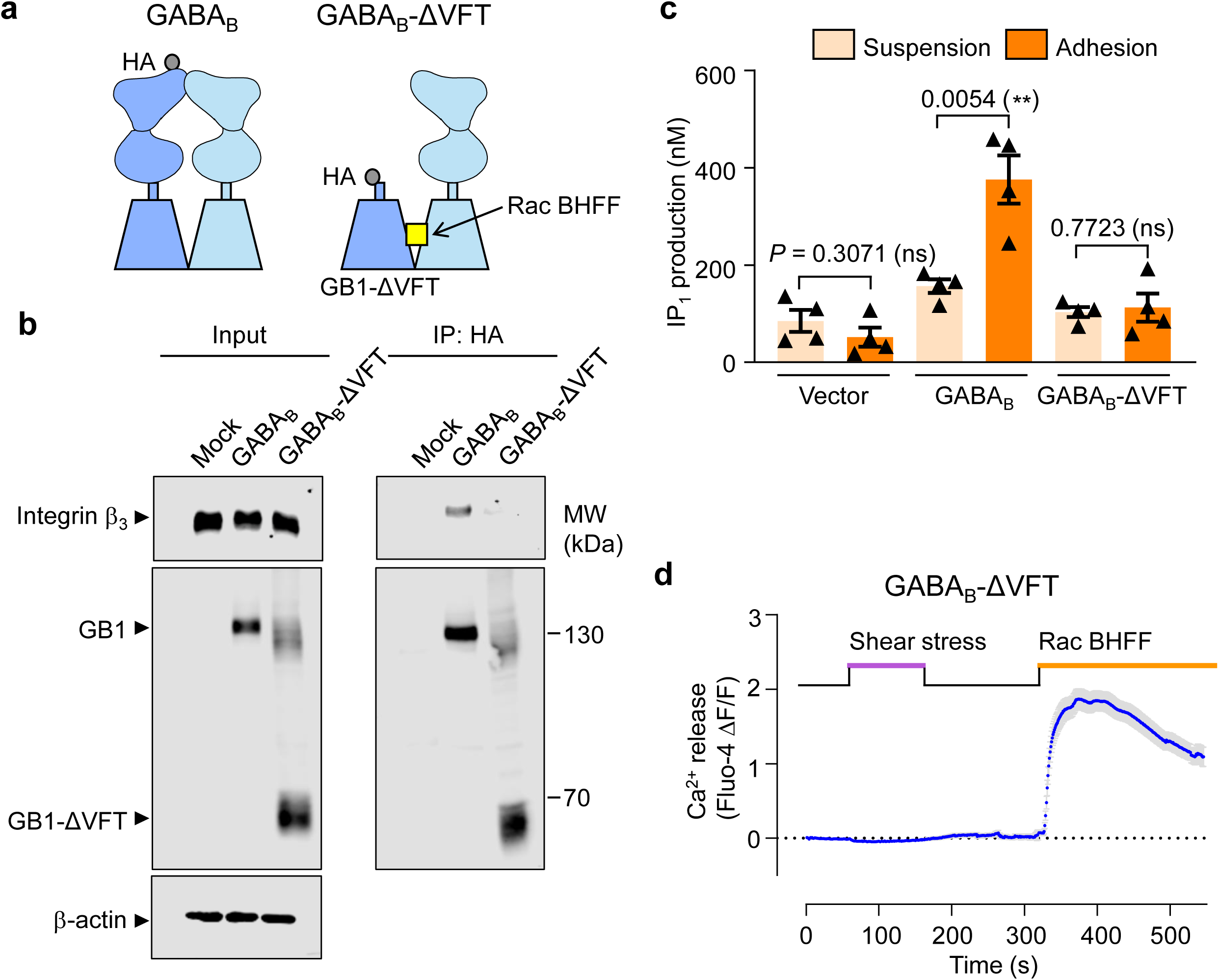
| VFT of GB1 is responsible for the interaction between GABA_B_ receptor and integrin β_3_, and the mechano-activation of GABA_B_ receptor. **a**, Schematic representation of GABA_B_-ΔVFT truncation, in which GB1 subunit lacks of the VFT domain (GB1-ΔVFT), but retains the ability to be activated by positive allosteric modulator Rac BHFF (yellow square). Co-immunoprecipitation experiments were performed using anti-HA antibody targeting to the HA tag, which is fused in the N-terminal of GB1 or GB1-ΔVFT. **b,** Co-immunoprecipitation of GABA_B_ receptor or GABA_B_-ΔVFT and integrin β_3_ using anti-HA antibody. Blots are from one representative of five biologically independent experiments. **c,** IP_1_ production in cells transfected with GABA_B_ receptor, or GABA_B_-ΔVFT, along with G_qi9_ under suspension or adhesion conditions. Data are present as mean ± s.e.m. from four biologically independent experiments each performed in triplicates and analyzed using unpaired t test (two-tailed) to determine significance. \**P* < 0.05, not significant (ns) > 0.05. **d,** Real-time recording of intracellular Ca^2+^ release in HEK293 cells expressing GABA_B_-ΔVFT and G_qi9_. After recording the basal state of Ca^2+^ release for 50 seconds, cells were subjected to shear stress for 100 seconds. Shear stress was then halted for 150 seconds, after which Rac BHFF was added for 200 seconds. Data are present as mean ± s.e.m. from 135 cells recorded.

Next, we used a glycan wedge scanning approach to map the interaction interface between the VFT_GB1_ and integrin β_3_. A consensus sequence NXS/T was introduced into GB1_VFT_ to enable the conjugation of a bulky N-glycan moiety to the side chain of the Asn residue, thereby forming a steric wedge to prevent GB1 from interacting with other proteins^66^. We selected seven GB1 mutants: 320^STL^322 (M1), 326^EER^328 (M2), 330^KEA^332 (M3), 337^TFR^339 (M4), 347^AVP^349 (M5), 352^NLK^354 (M6) and 356^QDA^358 (M7) (**Fig. 5a-c**). We have previously shown that the presence of an N-glycan moiety in these mutants do not affect agonist-induced GABA_B_ receptor activation^66^. We found that although the traction force failed to activate M2, M4, M6 and M7, it was still able to activate M1, M3 and M5 (**Fig. 5d**). Moreover, the amount of integrin β_3_ co-IPed with GB1 was greatly reduced in M7 mutant, but not in M3 mutant (**Supplementary Fig. 4b**). This indicates that residues 326-328, 337-339, 351-353 and 356-358 in GB1_VFT_ are important for its interaction with integrin β_3_. Interestingly, these sequences in GB1 are highly conserved across different species (*human*, *mouse*, *C. elegant*, *D. melanogaster*, *R. norvegicus*, *B. taurus*, *C. sabaeus*), especially the last 5 residues RQDAR (**Supplementary Fig. 4c**). We thus tested this RQDAR motif in GB1_VFT_, and found that the GABA_B_-5A mutant (GB1^RQDAR-5A^ + GB2) (**Fig. 5b**), in which RQDAR were mutated into five A (Ala) residues, exhibited a much weaker interaction with integrin β_3_ (**Supplementary Fig. 4d**). Importantly, this GABA_B_-5A mutant could still be activated by GABA (**Fig. 5e**), but lost sensitivity to traction force and shear stress **(Fig. 5f-g**). This set of experiments demonstrate that the GB1 subunit of GABA_B_ receptor interacts with integrin β_3_ via its VFT domain and that the RQDAR motif in VFT_GB1_ is important for GABA_B_ receptor’s interaction with integrin β_3_ as well as its activation by mechanical forces.

**Fig. 5.**
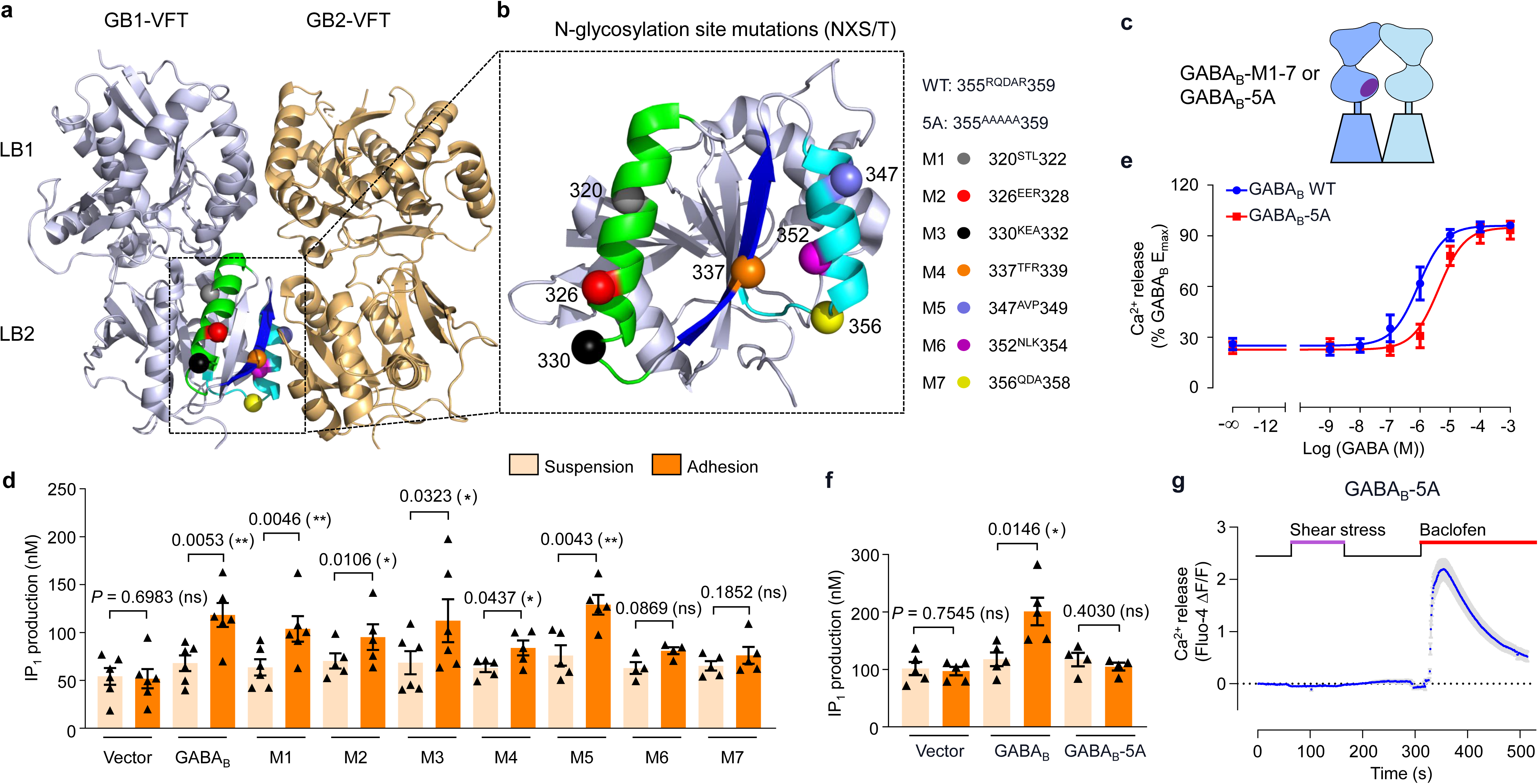
| Mapping the interaction region between GABA_B_ receptor and integrin β_3_. **a**, 3D model of the VFT of GB1 and GB2. N-glycosylation mutations (M1-7) are highlighted in LB2 of GB1 VFT. **b,** Detailed presentation of the mutants of M1-7 and 5A. **c,** Schematic representation of GABA_B_-M1-7 and GABA_B_-5A. **d,** IP_1_ production in HEK293 cells transfected with GABA_B_ receptor WT, or GABA_B_ receptor mutants M1-7 along with G_qi9_ under suspension or adhesion conditions. Data are present as mean ± s.e.m. from at least three biologically independent experiments (from left to right: n = 6, 6, 6, 5, 6, 5, 5, 4, 5) each performed in triplicates and analyzed using paired t test (two-tailed) to determine significance. \*\**P* < 0.01, \**P* < 0.05, not significant (ns) > 0.05. **e,** Intracellular calcium release induced by different dose of GABA in HEK293 cells transfected with GABA_B_ receptor WT and G_qi9_, or GABA_B_-5A and G_qi9_. Data are presented as mean ± s.e.m. from three biologically independent experiments each performed in triplicates. **f,** IP_1_ production in HEK293 cells transfected with GABA_B_ receptor WT and G_qi9_, or GABA_B_-5A and G_qi9_ under suspension or adhesion conditions. Data are mean ± s.e.m. from at least four biologically independent experiments each performed in triplicates and analyzed using unpaired t test (two-tailed) to determine significance. \**P* < 0.05, not significant (ns) > 0.05. **g,** Real-time recording of intracellular Ca^2+^ release in HEK293 cells expressing GABA_B_-5A and G_qi9_. After recording the basal state of Ca^2+^ release for 50 seconds, cells were subjected to shear stress for 100 seconds. Shear stress was then halted for 150 seconds, after which baclofen was added for 200 seconds. Data are present as mean ± s.e.m. from 111 cells recorded.

### Mechano-force acts as a positive allosteric modulator agonist (PAM ago) for GABA_B_ receptor activation

Ligand-induced GABA_B_ receptor activation features a close contact between LB2_GB1_ and LB2_GB2_, further triggers an allosteric rearrangement of 7TMs at the TM6-TM6 interface between GB1 and GB2 subunits (**Fig. 6a**), which is considered a hallmark of the active state of GABA_B_ receptor^30, 32–35^. Upon ligand binding, the two Val residues in TM6 of GB1^6.56^ and GB2^6.56^ turn close to each other in the active state of the receptor^35^ (**Fig. 6b**). The close proximity of the two Val residues in this active state enables the formation of GB1-GB2 dimers through covalent cross-linking of the two subunits, in which the two Val residues are mutated to Cys residue: GB1^6.56C^ and GB2^6.56C^ ^35^. Thus, this assay allows the probing of the active state of GABA_B_ receptor in HEK293 cells transfected only with GB1^6.56C^ and GB2^6.56C^. Consistent with our previous reports, GABA treatment promoted the formation of GB1^6.56C^-GB2^6.56C^ dimers^35^. Of primary significance, shear stress also promoted the formation of GB1^6.56C^-GB2^6.56C^ dimers (**Fig. 6c**). Furthermore, FN-treatment also increases of formation of GB1^6.56C^-GB2^6.56C^ dimers (**Fig. 6d**). Finally, we used a FRET-based inter-subunit sensor of the GABA_B_ receptor to measure the conformational change between two VFTs of GABA_B_ receptor as previously reported by us^67^ (**Fig. 6e**). The FRET ratio was increased under FN treatment (**Fig. 6f**), suggesting that interaction between FN and integrin induce a change in the conformation of two VFTs to bring the GB1^LB2^ and GB2^LB2^ in contact. In all, these set of experiments demonstrate that mechanical forces can facilitate an allosteric interaction between two VFTs to further induce re-arrangement of 7TMs of GABA_B_ receptor towards an active conformation in a manner similar to that induced by ligand binding, demonstrating that the mechanic force acts as a PAM ago for GABA_B_ receptor activation.

**Fig. 6.**
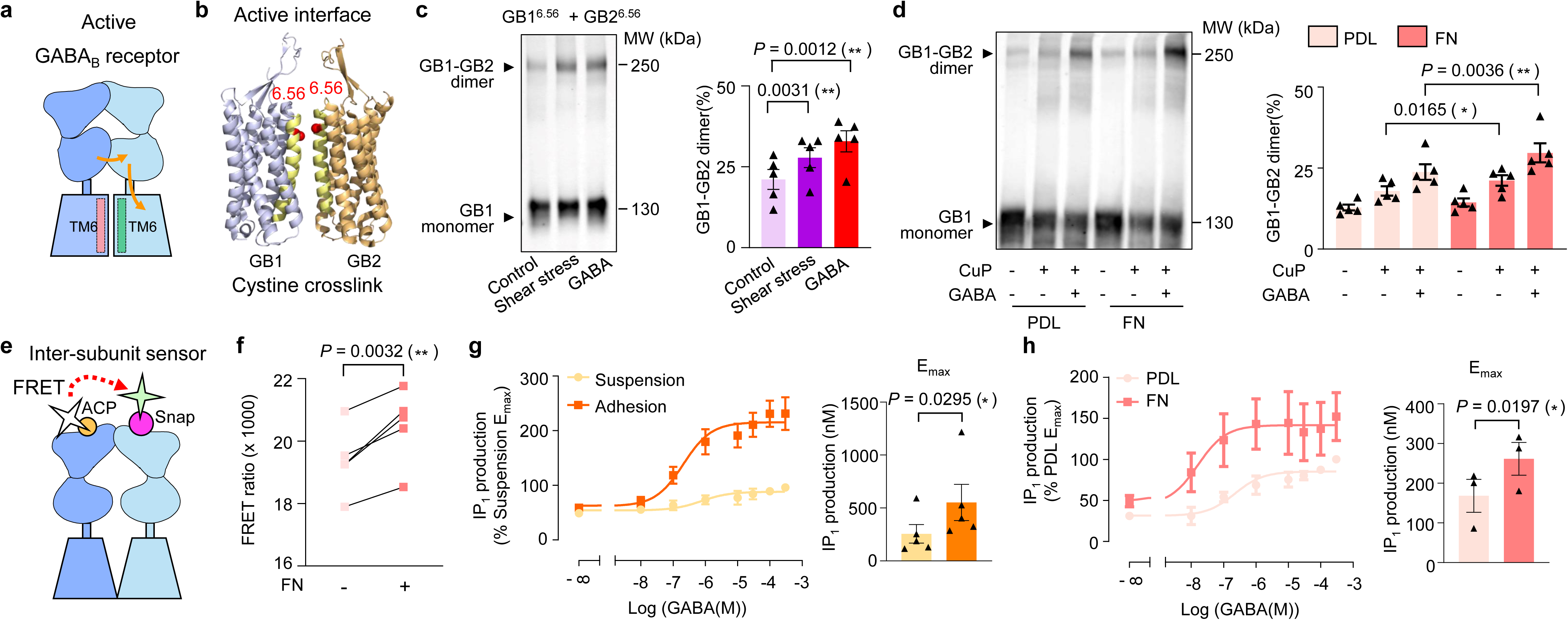
| Mechano-force acts as a positive allosteric modulator agonist (PAM ago) for GABA_B_ receptor activation. **a**, Schematic representation of the active state of GABA_B_ receptor, while the TM6s of GB1 and GB2 are rearranged towards each other. **b,** Highlight of the cysteine substitutions (red balls) in the position 6.56 of TM6 in GB1 and GB2 in its active state structure (PDB: 7EB2). **c,** Cross-linking of cell surface GABA_B_ receptor between the two cysteines in GB1^6.56^ and GB2^6.56^ with CuP treatment along with no shear stress (control), shear stress treatment (15 dyn/cm^2^, 30 min) or GABA treatment (100 μ, 30 min). **d,** Cross-linking of cell surface GABA_B_ receptor dimer between the two cysteines in GB1^6.56^ and GB2^6.56^ encompassing three conditions: without CuP treatment, with CuP treatment, with both CuP and GABA treatment, under PDL or FN-treated condition. MW, molecular weight. Blots in *(c-d)* are from one representative experiment of five biologically independent experiments. The bars show the percentage of GB1-GB2 dimer, which are relative to total amount of GB1 including GB1-GB1 dimer and GB1 monomer in each lane. Data are present as mean ± s.e.m. from five biologically independent experiments respectively in *(c-d)* and analyzed using paired t test (two-tailed) to determine significance. \**P* < 0.05, \*\**P* < 0.01. **e,** Schematic representation of the inter-subunit sensor of GABA_B_ receptor, measuring the conformational change between VFT_GB1_ and VFT_GB2_. The LB1_GB1_ is tagged with ACP and labeled with fluorescent molecular as energy donor, while LB1_GB2_ is tagged with Snap and labeled with fluorescent molecular as energy acceptor. FRET can be measured between two fluorescent molecules. **f,** FRET ratio measured using the inter subunit sensor of GABA_B_ receptor shown in *(e)*. HEK293 cells were transfected with GABA_B_ sensors and cultured under PDL and FN coating condition. Data are present as mean ± s.e.m. from five biologically independent experiments each performed in triplicates and analyzed using paired t test (two-tailed) to determine significance. \*\**P* < 0.01. **g-h,** Dose response curve and E_max_ analysis of GABA-induced IP_1_ production in HEK293 cells transfected with GABA_B_ receptor and G_qi9_ under suspension or adhesion conditions *(g)*, or under PDL or FN-treatment *(h)*. Data are present as as mean ± s.e.m. from five and three biologically independent experiments each performed in triplicates in *(g)* and *(h)* respectively. E_max_ are analyzed using paired t test (two-tailed) to determine significance. \**P* < 0.05.

More interestingly, we also observed that FN-treatment also increases of GABA-induced formation of GB1^6.56C^-GB2^6.56C^ dimers **(Fig. 6d**), suggesting the PAM effect of mechano-force on GABA-induced activation of GABAB receptor. We further verified whether and how mechanical forces allosterically regulate GABA-induced GABA_B_ receptor activity. Interestingly, both the efficacy and potency of GABA-induced receptor activation was increased under adhesion and FN-treated conditions (suspension vs adhesion, E_max_: 256 nM vs 552 nM, EC_50_: 0.18 μM vs 0.08 μM; PDL vs FN, E_max_: 168 nM vs 257 nM, EC_50_: 0.19 μM vs 0.02 μM) (**Fig. 6g-h**), highlighting the potential PAM effect of mechano-force on GABA-induced GABA_B_ receptor function *in vivo*.

### GABA_B_ receptor is required for shear stress-induced remodeling of astrocytes

Having characterized the mechanisms underlying GABA_B_ receptor activation by mechanical forces, we investigated whether and how mechanical force may regulate cellular physiology. GABA_B_ receptors are widely expressed in many types of neurons in the brain^8^. However, the relative low abundance and high heterogeneity of neurons (compared to glial cells) pose a challenge for investigating this question in neurons. In addition to neurons, GABA_B_ receptors are also broadly expressed in glial cells, particularly in astrocytes^9, 11^, which are the most abundant neural cells in the brain^68^. In response to mechanical insults experienced during traumatic brain injury, astrocytes undergo morphological, molecular and functional remodeling, which transforms them into reactive astrocytes in a process called reactive astrogliosis^69^. Reactive astrocytes play an important role in tissue repair and remodeling following brain trauma^70–72^. Other pathological conditions such as infection, ischemia, epilepsy, and cancer also trigger reactive astrogliosis^71, 73–76^. The robust remodeling of astrocytes induced by mechanical insults, high abundance of these glial cells in the brain, and relative ease of culture prompted us to explore whether GABA_B_ receptors contribute to mechano-induced reactive astrogliosis. To do so, we cultured primary astrocytes isolated from the mouse brain and assayed the effects of shear stress on their remodeling. We found that shear stress greatly increased the size of astrocytes and their expression of glial fibrillary acidic protein (GFAP), two key parameters of astrocyte reactivity^76, 77^, indicating that shear stress promoted reactive astrogliosis (**Fig. 7a-c**). Importantly, siRNA knockdown of the GABA_B_ receptor abolished the shear stress-induced increase in the size of astrocytes as well as the expression level of GFAP, pointing to a key role of GABA_B_ receptor in the process (**Fig. 7a-d**). Additionally, baclofen increased both cell size and GFAP expression of astrocytes, which mimicked the effect of shear stress, demonstrating the physiological role of GABA_B_ receptor activation in astrocyte remodeling (**Supplementary Fig. 5**). Together, these results demonstrate that shear stress-induced reactive astrogliosis requires GABA_B_ receptors.

**Fig. 7.**
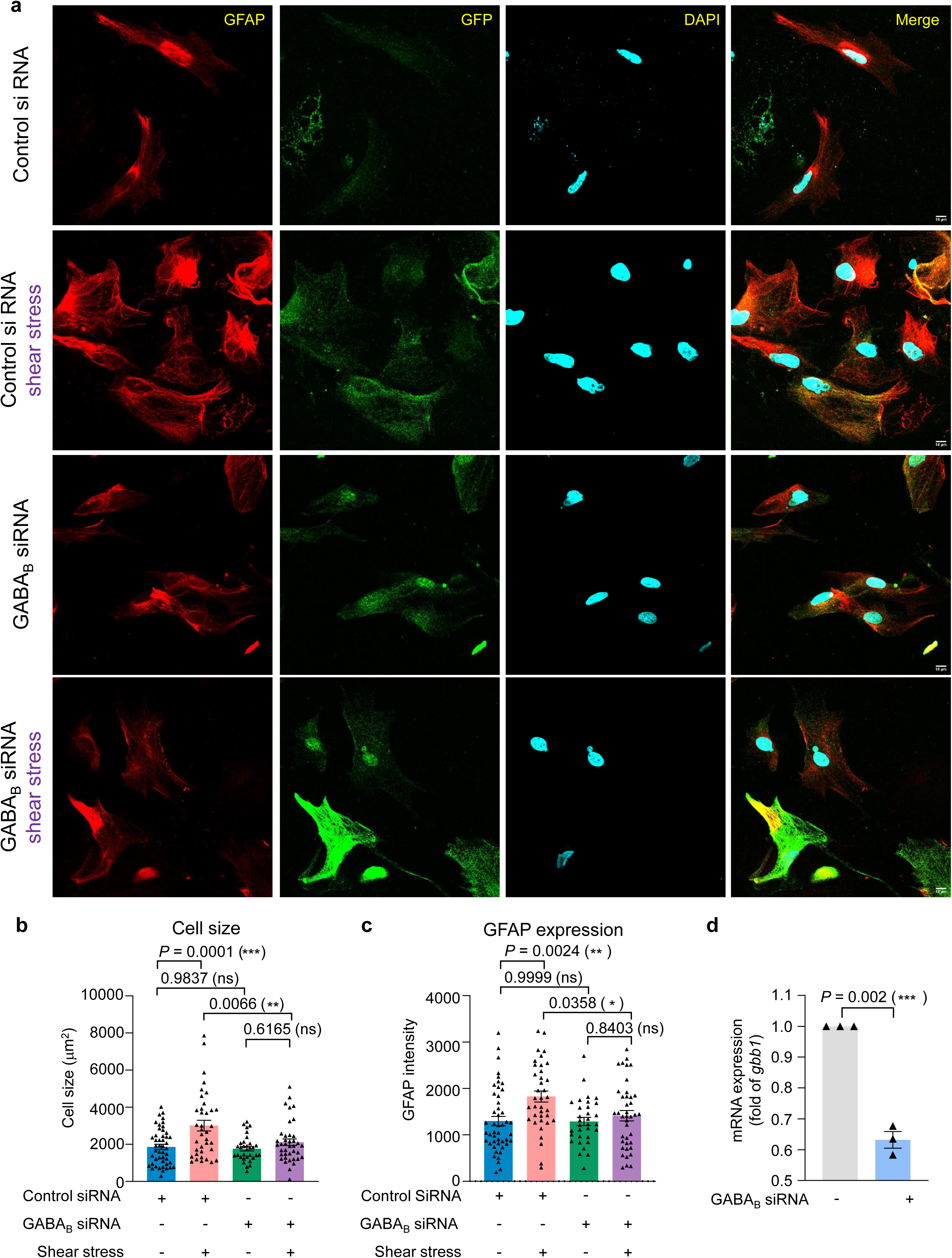
| GABA_B_ receptor is required for shear stress-induced astrocyte remodeling. **a**, Immunofluorescent staining of GFAP (red), GFP (green) and DAPI (cyan) in astrocytes transfected with control siRNA or GABA_B_ receptor siRNA along with GFP, treatment with or without shear stress (15 dyn/cm^2^, 30 min). Images are representative from three biologically independent experiments. Scale bar: 10 μm. **b-c,** Analysis of the cell size and GFAP expression of astrocytes with the same treatment in *(a)*. Measurements are made on each cell by cell basis (ROI) from three biologically independent experiments (number of cells from left to right: n = 47, 37, 32, 39). Data are present as mean ± s.e.m and analyzed using ordinary one-way ANOVA with Tukey’s multiple comparisons test to determine significance. \*\*\**P* < 0.001, \*\**P* < 0.01, ns: non-significant. **d,** RNA interference efficacy of the GB1 in astrocytes in *(b-c)* using qPCR detecting *gbb1* expression. Data are present as mean ± s.e.m. from three biologically independent experiments and analyzed with unpaired t test (two-tailed) to determine significance. \*\*\**P* < 0.001.

### Shear stress activates astrocytes in a GABA_B_ receptor-dependent manner

The finding that GABA_B_ receptors are required for shear stress to promote reactive astrogliosis suggests that shear stress may activate GABA_B_ receptor in these primary cells just like when it was expressed in HEK293 cells. To test this hypothesis, astrocytes were recorded using calcium imaging to determine their response to shear stress. In astrocytes, Ca^2+^ is a major downstream event of GABA_B_ receptor and dependent on G_i/o_ protein, as previously reported^9, 78–81^. Therefore, no transfection of G protein chimera was required to detect GABA_B_ receptor-induced Ca^2+^ signal in astrocytes. The cells with Ca^2+^ activation was characterized as previously^82^ by an increasing in the Ca^2+^ signal to reach a peak after stimulus and sustain for more than 5s. Owning to the heterogeneity of primary astrocytes, approximately 37.29% of astrocytes were sensitive to baclofen and were found to induce Ca^2+^ release, similar to a previous report^9^. About half (45.76%) of the astrocytes were sensitive to shear stress with a transient but significant increase of the Ca^2+^ signal (**Fig. 8a**). Among them, 62.04% were sensitive to baclofen, a GABA_B_ receptor-specific agonist, indicating that most shear stress-sensitive astrocytes expressed GABA_B_ receptors (**Fig. 8a-b**). Notably, siRNA knockdown of the GABA_B_ receptor greatly reduced the population of astrocytes that were sensitive to baclofen (reduced to 6.69% from 37.29%) (**Supplementary Fig. 6a-c**) as well as shear stress (reduced to 13.48% from 45.76%) (**Fig. 8c-d**), indicating that GABA_B_ receptors play a crucial role in mediating shear stress-induced response in these primary cells. The strength of the Ca^2+^ signal was significantly decreased for both shear stress and baclofen treatment, when GABA_B_ receptor was knocked down, whereas as a control, the population of astrocytes with ATP sensitivity remained 100% and the Ca^2+^ signal was still strongly activated (**Supplementary Fig. 6d**). Overall, we demonstrated that shear stress can activate astrocytes in a GABA_B_ receptor-dependent manner. This suggests a model in which shear stress can activate GABA_B_ receptor in astrocytes, triggering reactive astrogliosis.

**Fig. 8.**
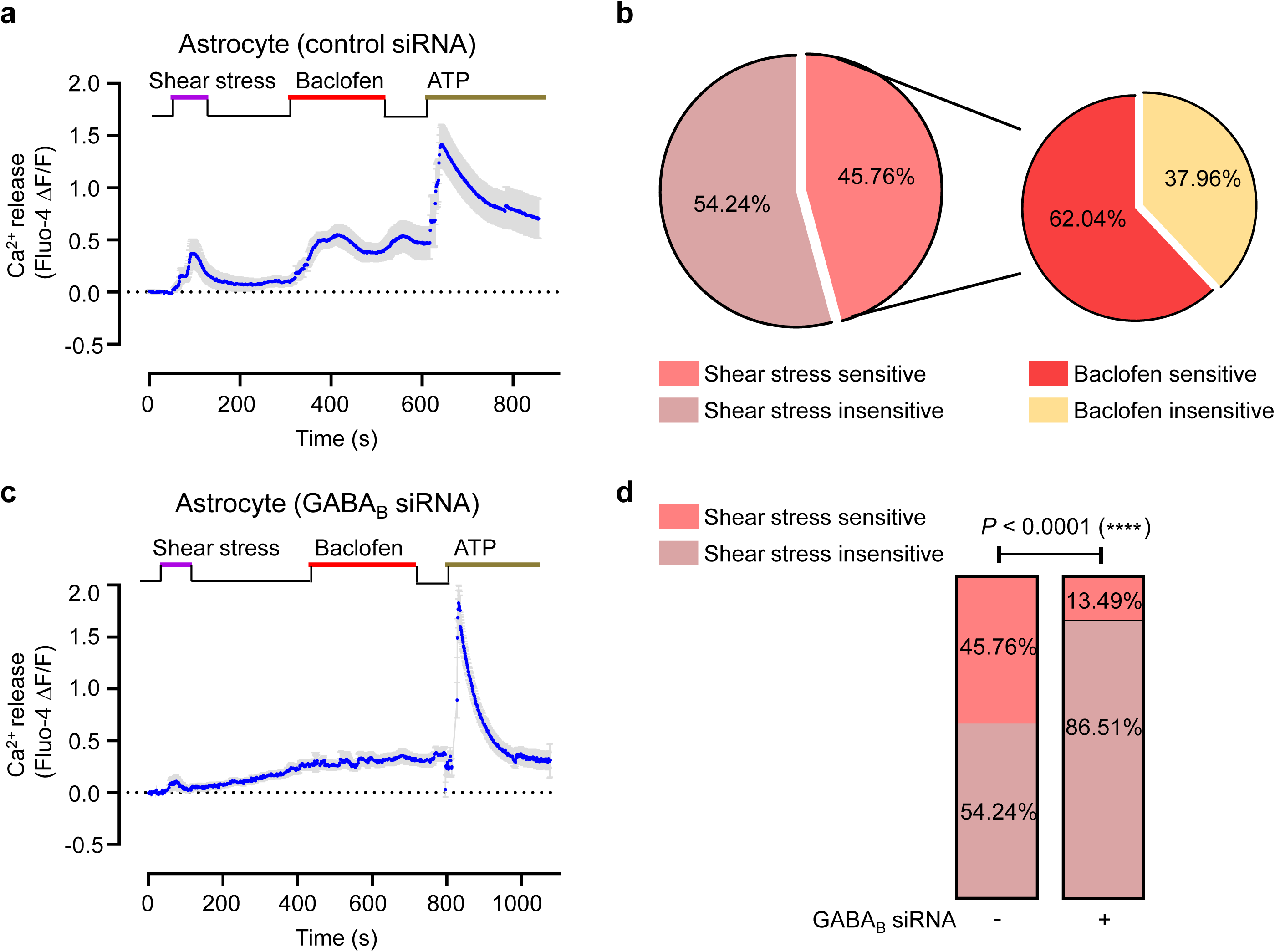
| Shear stress activates GABA_B_ receptor in astrocytes. **a**, Real-time recording of intracellular Ca^2+^ release in astrocytes under shear stress. Cells were transfected with control siRNA and treated with shear stress, baclofen (100 μM) or ATP (100 μM) in the indicated time point. Data are representative from six biologically independent experiments. **b,** Percentage of astrocytes in response to shear stress or baclofen in 236 recorded cells. **c,** Real-time recording of intracellular Ca^2+^ release in astrocytes under shear stress. Cells are transfected with siRNA knocking down GABA_B_ receptor and administrated to shear stress, baclofen (100 μM) or ATP (100 μM) in the indicated time point. Data are representative from six biologically independent experiments. **d,** Percentage of astrocytes with the calcium response (sensitive) or without the calcium response (insensitive) to shear stress (control siRNA: 236 cells; GABA_B_ siRNA: 314 cells). Data are analyzed with χ2 test to determine significance. \*\*\*\**P* < 0.0001.

## Discussion

As a classic class C GPCR, the activation mechanism of GABA_B_ receptor has been extensively characterized at both the biochemical and structural levels, which features a series of ligand binding-induced structural rearrangements in the receptor, culminating in its formation of a ternary complex with heterotrimeric G_i/o_ proteins^30, 83^. In this study, we found that GABA_B_ receptors were activated by mechanical forces in a GABA-independent manner. To the best of our knowledge, this is the first report of a ligand-independent activation mechanism for GABA_B_ receptor.

It is notable that the GABA_B_ receptor forms a protein complex with integrin and that GABA_B_ receptor sensing of mechanical forces requires a direct interaction with integrin. We thus propose that GABA_B_ receptor and integrin form a mechano-transduction complex. In this complex, integrin likely functions as a mechano-sensor, as integrin is well-known to sense mechanical forces such as traction force and shear stress in cellular mechanotransduction^51, 84^. On the other hand, as blocking integrin function or inhibiting its expression abolishes the mechano-activation of GABA_B_ receptors, GABA_B_ receptors may not directly sense mechanical forces but rather function as a transducer that transduces mechanical signals to downstream G proteins. Here we showed that GABA_B_ receptor interacts with the integrin subunit β3 and β1. Other integrins may also be involved as FN and RGDS can bind to several integrin types^55, 60^. A growing list of GPCRs have been reported to be mechanosensitive, such as AT1R^85, 86^, H1R^86, 87^, bradykinin B2R^88^, GPR68^89^, and adhesion GPCRs^90–92^. Unlike GABA_B_ receptors, these mechanosensitive GPCRs are believed to directly sense mechanical forces as a mechano-sensor, though the underlying mechano-transduction mechanisms are not well understood. Thus, GABA_B_ receptor represents the first GPCR that acts as a mechano-transducer rather than a mechano-sensor, revealing a novel role for GPCRs in mechano-transduction.

Strikingly, although mechanical forces activate GABA_B_ receptor in the absence of ligand binding, they trigger an allosteric rearrangement of the 7TMs in GABA_B_ receptor towards an active state in a manner similar to that induced by ligand binding. Specifically, both ligand binding and shear stress induced a similar allosteric rearrangement of 7TMs in the TM6-TM6 interface of the GB1 and GB2 subunits. This structural rearrangement has been shown to open a shallow cavity at the intracellular side of GABA_B_ receptor to provide access for G protein binding^30, 32–35^, which is a hallmark of the active state of GABA_B_ receptor. Notably, agonists such as GABA and baclofen bind to a larger crevice between LB1 and LB2 of VFT_GB1_ to bring the two LB2 of two subunits in contact. As shear stress promotes the binding of integrin to LB2 of VFT_GB1_, it is conceivable that shear stress may activate the GABA_B_ receptor by promoting integrin binding to LB2 of VFT_GB1_, which in turn pushes the two LB2 in contact and triggers an allosteric rearrangement of 7TMs in the TM6-TM6 interface towards an active conformation, culminating in receptor activation. In this regard, shear stress-induced binding of integrin to GB1 would mimic the role of ligand (e.g. GABA) binding to GB1 in GABA_B_ receptor activation, providing a potential molecular mechanism by which mechanical forces activate GABA_B_ receptor and allosterically potentiate the GABA effect as a PAM ago (**Fig. 9**). However, though both GABA and mechanical force lead to GABA_B_-mediated G activation, the mechanism is not exactly the same. The FRET between the N-terminal of GB1 and GB2 subunit showed higher signal under mechanical force, while GABA induced a decreased FRET signal. These two observations are not contradictory, because the closure of GB1 VFT is the critical step to initiate the receptor activation. Further structural analysis of GABA_B_ receptor and integrin complex will be needed to fully understand the activation mechanism. It is also different from the adhesion GPCRs, which act as mechano-sensors and rely on an extremely large N terminus for ECM recognition and an autoproteolysis domain that undergoes self-cleavage for receptor activation^92, 93^. The GABA_B_ activation displayed faster under baclofen treatment than shear stress induced Ca^2+^ signal in our experiments. This faster chemical response than mechno-force was also observed in GPR68^89^. However, it doesn’t indicate a faster response to baclofen than to shear stress, as these two stimuli are not directly comparable. A saturating concentration of baclofen can be applied to fully activate the GABA_B_ receptor, whereas higher shear stress speeds would detach cells, preventing reliable signal detection. This fundamental difference precludes a strict kinetic comparison.

**Fig. 9.**
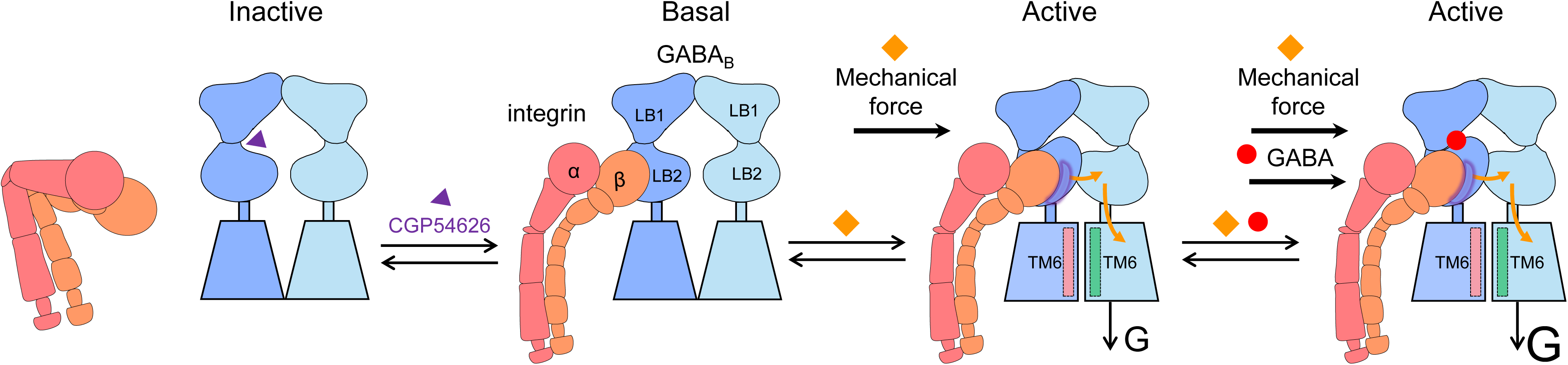
| Schematic model of GABA-independent allosteric activation of GABA_B_ receptor by mechanical forces through integrin interaction. Antagonist CGP54626-bound GABA_B_ receptor fails to interact with integrin β3 whereas integrin β3 interacts with GB1_VFT_ of GABA_B_ receptor with constitutive activity. Mechanical force promotes GB1_VFT_ and integrin β3 interaction to stabilize the closure of GB1_VFT_-induced LB2_GB1_ and LB2_GB2_ in contact, further inducing an allosteric re-arrangement of the GABA_B_ receptor 7TM towards an TM6-TM6 active conformation, culminating in the asymmetric activation of the receptor with G protein under GB2. Mechanical force acts as a positive allosteric modulator to boost up GABA-induced GABA_B_ receptor activation through interaction between GB1_VFT_ and integrin β3.

We showed that shear stress could stimulate the activity of primary astrocytes in a GABA_B_ receptor-dependent, but GABA-independent manner. We also found that shear stress promotes reactive astrogliosis, a process by which astrocytes undergo remodeling in response to pathological conditions, such as brain trauma, to transition into reactive astrocytes that are important for tissue repair^70–72^. Although the brain is typically considered as a protected organ, it may also experience transient shear stress, such as during traumatic brain injury, which can generate both localized and distributed forces throughout the brain^70, 71, 94^. Importantly, shear stress-induced reactive astrogliosis requires GABA_B_ receptor. These results together suggest a model in which mechanical forces activate GABA_B_ receptors in astrocytes to promote reactive astrogliosis. Though we confirmed no GABA in our experiments, the endogenous GABA is present in vivo, which may work synergistically together with shear stress. Although GABA_B_ receptor appears to a major contributor to astrocyte mechano-sensitivity, siRNA knockdown of the GABA_B_ receptor eliminated most but not all mechanosensitive astrocytes. This is consistent with the notion that additional mechano-sensors are present in these glial cells, such as mechanosensitive Piezo channels as reported recently^95^.

As the sole metabotropic GABA receptor, GABA_B_ receptor regulates a wide spectrum of physiological processes, ranging from synaptic inhibition to locomotion^7^, cognition^6^, addiction^18^, epilepsy^12, 13^, and astrocyte morphogenesis^10, 11^. To date, all the physiological functions carried out by GABA_B_ receptor have been attributed to its activation by GABA. Our discovery of a GABA-independent activation mechanism of the GABA_B_ receptor and its crucial role in promoting astrocyte remodeling raises the possibility that this novel mode of activation may contribute to additional aspects of GABA_B_ receptor functions in the brain. Our work also raises the possibility that other GPCRs could be potentially sensitive to mechanical forces by forming a mechano-transduction complex with integrin.

## Materials and methods

### Antibodies

Mouse monoclonal GB1 antibody targeting to the extracellular domain (#55051) was acquired from Abcam (Shanghai, China), and GB1 polyclonal antibody (#BS2717) was obtained from Bioworld Technology (Shanghai, China). Integrin β_3_ antibody (#sc-46655) and anti-mouse IgG (#sc-2025) for co-immunoprecipitation experiment were purchased from Santa Cruz Biotechnology (Shanghai, China). Integrin α_v_ antibody (#A2091) was purchased from ABclonal Technology (Wuhan, China). Integrin β_3_ antibody (#13166), β-actin (#4967), phospho-Myosin Light Chain 2 (MLC-PP, #3675), GFAP (#3670), anti-mouse IgG HRP-linked antibody (#7076), anti-rabbit IgG HRP-linked antibody (#7074), anti-rabbit IgG (H+L) (DyLight^TM^ 800 4X PEG Conjugate) (#5151) and anti-mouse IgG (H+L) (DyLight^TM^ 800 4X PEG Conjugate) (#5257) were purchased from Cell Signaling Technology (Shanghai, China). Anti-HA antibody (#3F10) was purchased from Roche (Shanghai, China). Cy3 AffiniPure Donkey Anti-Mouse IgG (H+L) (A0521) was purchased from Beyotime Biotechnology (Shanghai, China).

### Reagents

Dulbecco’s Modified Eagle Medium (DMEM, #8120251), fetal bovine serum (FBS, #10437028), penicillin-streptomycin (#15140-122), Opti-MEM medium (#31985070), cell dissociation buffer (#2075828), Fluo4-AM (#F14202), lipofectamine 2000 (#11668019) and Pierce^TM^ enhanced chemiluminescence reagents (#37074) were purchased from Thermo Fisher Scientific (Shanghai, China). Proteinase cocktail inhibitor (#04693159001) was bought from Roche (Shanghai, China). Protein G beads (#3394201) and nitrocellulose membranes (#HATF00010) were from Millipore (Shanghai, China). GABA (γ-aminobutyric acid) (#43811), Dichloro (1, 10-phenanthroline) copper (II) (#362204), Poly-D-Lysine (#P1149) and Poly-L-ornithine hydromide (#P3655) were bought from Sigma (Shanghai, China). Blebbistatin (#B1387) was purchased from ApexBio Technology (Shanghai, China). Recombinant Human Fibronectin (#40113ES10) was from Yeasen (Shanghai, China). CGP54626 hydrochloride (#HY-101378) and RGDS peptide (Arg-Gly-Asp-Ser, #HY-12290) were from Med Chem Express (Shanghai, China). Baclofen (#0796) and Rac BHFF (#3313) were from Tocris Bioscience (Shanghai, China). Coelenterazine H (#S2011) was purchased from Promega (Beijing, China). Snap-Surface Alexa Fluor 647 (#S9136) was from New England Biolabs (Ipswich, MA, USA).

### Plasmids

The pRK5 plasmids encoding N-terminal HA-tagged wild-type rat GB1a, GB1^ASA^, GB1^Δ^_VFT_, N-terminal Flag-tagged wild-type rat GB2 were described previously^36, 66^. ^HA-Snap-^GB1, ^HA-Snap-^GB1^6.56C^, and ^Flag-Halo-^GB2^6.56C^ have been reported previously^35^. The pRK5 plasmids encodes the GB1^ΔVFT^, tagged with HA and Snap inserted after the signal peptide. The pRK5 plasmids encoding wild-type human Integrin β_3_ (UniProt: P05106) was tagged with HA tag, inserted immediately after the signal peptide. The Integrin α_v_ plasmid (UniProt: P06756, #P40612) was purchased from the company (Miaolingbio, Wuhan, China). The probes (full-length mVenus, Rluc, YFP or RFP) were fused to the C terminus of the GB1, GB2, Integrin β_3_, Integrin β_1_, Integrin α_v_, mGlu2 or 5-HT_2a_ receptors. G_i_ protein sensors including G_i1_-Nluc, G_β1_ and G_γ2_-Venus were previously described^49^. All the mutants for GABA_B_ receptor were generated by site-directed mutagenesis using the QuikChange mutagenesis protocol (Agilent Technologies, Stratagene, La Jolla, CA).

### HEK293 Cell culture and transfection

HEK293 cells (ATCC, CRL-1573, lot: 3449904) were cultured in DMEM supplemented with 10% fetal bovine serum (heat-inactivated) at 37 °C under 5% CO_2_. Transfection was performed using lipofectamine 2000 following manufacturer protocol. Two million cells were transfected with indicated plasmids with the lipofectamine/DNA ratio at 2:1, and the total DNA at 1.5 μg in 35 mm diameter dishes.

### Primary astrocyte culture

All experiments were approved by the Animal Experimentation Ethics Committee of School of Life Science and Technology at Huazhong University of Science and Technology and were specifically designed to minimize the number of animals used. Primary astrocytes were cultured as described previously^96^. Briefly, the cerebral cortex was dissected from KunMing mice (one day old) obtained from the Hubei Provincial Center for Disease Control and Prevention. Following careful removal of meninges, tissues were dissected and cut into small pieces using sharp blades, then digested in 1-2 ml of Trypsin/EDTA in a 37 °C water bath for 8 min. The supernatant was discarded, and the tissue was gently triturated using a 1 ml pipette. The homogenate was centrifuged at 180 g for 5 min. The pellet was re-suspended and seeded in a T75 flask pre-coated with poly-L-ornithine. Cells were cultured in DMEM medium, supplemented with 10% FBS at the density of 1 × 10^4^/cm^2^. 24 hours after the initial plating, the media were changed with DMEM supplemented with 10% FBS, 100 units/mL of penicillin, and 100 µg/mL of streptomycin and suspended cells were removed. The cells were cultured at 37 °C in a 5% CO_2_ incubator for 7 days. The dishes were shaken at 260 rpm (2 h, 37 °C) to obtain purified astrocytes. The remaining enriched astrocyte culture was exposed to 0.5% trypsin for 5 min to cause detachment of the glial monolayer and cells were then re-plated in a new T75 flask to further purify astrocytes. The astrocytes were used after three times of passage at day 21 for all the experiments.

### Cell suspension and adhesion experiment

Cell adhesion experiments were performed as previously described^97^. HEK293 cells were suspended using cell dissociation buffer and collected by centrifugation (180 g, 5 min) to remove the supernatant. Then cells were suspended in PBS and seeded into 96-well plate. For the suspension condition, the well was coated with 1% BSA. For the adhesion condition, the surface of the well was coated with poly-D-lysine (PDL) (5 μg/cm^2^). The cells were kept in the wells for 3-4 h at 37 °C under 5% CO_2_, then lysis buffer was added directly for further measurement of IP_1_ production measurements.

### PDL and FN treatment

Poly-D-lysine (PDL) coated surfaces were prepared by incubating 5 μg/cm^2^ PDL dissolved in phosphate-buffered saline in uncovered tissue culture dishes overnight at 37 °C. Fibronectin (FN)-coated surfaces were prepared by incubating 5 μg/cm^2^ FN diluted in phosphate-buffered saline (PBS) in tissue culture dishes overnight at 37 °C as reported^97^. The HEK293 cells expressing indicated plasmids were seeded into PDL-coated wells or FN-coated wells with culture medium for 24 h at 37 °C under 5% CO_2_. Then cells were lysed for IP_1_ production measurements.

### Shear Stress loading experiments

Shear stress was applied to cultured cells using an in vitro shear stress system as previous report^98^. A peristaltic pump (BT-CA600, Naturethink, China) was used to drive a stable flow shear stress. HEK293 cells were seeded on glass plates pre-coated with FN, then tranfected with indicated plasmids when cells grew to ∼80% confluence. After 24 h, glass plates were transferred onto a parallel-plate flow chamber (C901) and subjected to 15 dyn/cm^2^ (1.5 Pascals) of shear stress for 15 min, followed by IP_1_ production measurement.

### IP_1_ measurement

IP_1_ production was measured using the IP-One HTRF kit (62IPAPEB, Revvity, Codolet, France). Cryptate-labelled anti-IP_1_ monoclonal antibody and d2-labelled IP_1_ were diluted in lysis buffer and added to the cell lysis in 96-well plate. After 1 h of incubation at 25°C, the plates were read in PHERAstar FS (BMG Labtech, USA) with excitation at 337 nm and emission at 620 and 665 nm. The accumulation of IP1 concentration (nM) was calculated according to a standard dose response curve.

### G_i1_ protein dissociation measurement

G_i1_ protein dissociation measurements were performed as previously described^49^. HEK293 cells were transfected with Gi protein sensors including G_αi_-Nluc (1.5 ng), G_β1_ (10 ng) and Venus-G_γ2_ (10 ng), along with vector (control) or GB1 and GB2 (GABA_B_ receptor, 30 ng + 40 ng) per well in 96-well plate and cultured for 24h. Then cells were pre-incubated in PBS for 30 min. The signals emitted by the donor (460-500 nm band-pass filter, Em 480 nm) and the acceptor (510-550 nm band-pass filter, Em 530 nm) were recorded after the addition of 10 μM furimazine using the PHERAstar FS (BMG Labtech, USA) with the program PHERAstar control (Version 4.00 R4). All measurements were performed at 37°C. The BRET signal (BRET ratio) was determined by calculating the ratio between the emission of acceptor and donor (Em 530 nm / Em 480 nm).

### Laminar shear stress stimulation

Coverslips were plated in the laminar shear stress imaging chamber (open diamond bath imaging chambers RC-26, Warner) prior to experiments. Then chambers were mounted onto the microscope and connected to the peristaltic pump (ISM827B, Ismatec). Continuous laminar flow was applied to cells for 100 s.

Shear stress (τ) in arterial segments was calculated in the following equation, according to previous research^87^,

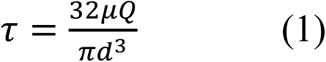

where Q is flow rate, μ is fluid viscosity and d is arterial diameter.

Polytetrafluoroethylene capillary tubes (d = 0.5 mm) were adapted to model arteries and only focused on the fluorescence from transfected cells next to the fluid input terminal, enabling to estimate the fluid shear stress on the surface of the coverslips. To calculate a specific value of shear stress, the flow rate (Q = 4.8 ml·min^-1^) of imaging buffer was controlled through a peristaltic-pump and measured the viscosity of fluid flow (μ = 0.681 ± 0.017 mPa·s, mean ± s.e.m., n = 8) through viscometers. Finally, all the parameters were bring into equation (1) to calculate the value of fluid shear stress (τ = 4.44 ± 0.11 Pa, mean ± s.e.m.).

### Calcium imaging

Calcium imaging was measured as previously reported^82^. For HEK293 cells, they were seeded onto FN-coated coverslips 6 hours after plasmid transfection. After another 24 hours, HEK293 cells were incubated with 1 μM Fluo4-AM in the imaging buffer (130 mM NaCl, 5.1 mM KCl, 0.42 mM KH_2_PO_4_, 0.32 mM Na_2_HPO_4_, 5.27 mM glucose, 20 mM HEPES, 3.3 mM Na_2_CO_3_, 1 mM MgSO_4_, 1.3 mM CaCl_2_, 0.1% BSA, 2.5 mM probenecid, pH adjusted to 7.4) for 30 min at 37°C. For primary astrocytes, they were incubated with 1 μM Fluo4-AM in the imaging buffer (145 mM NaCl, 2.5 mM KCl, 1 mM MgSO_4_, 2 mM CaCl_2_, 10 mM glucose, 10 mM HEPES, 3.3 mM Na_2_CO_3_, 1 mM MgSO_4_,

2 mM CaCl_2_, pH adjusted to 7.4) for 30 min at 37°C. Cells were washed with imaging buffer and pre-incubated for 5 min at room temperature before calcium imaging. Intracellular Ca^2+^ fluorescence intensity was measured at 485 nm to indicate basal signal (F_basal_) and response signal under stimulation of shear stress or baclofen (F_response_). The fluorescence ratio (ΔF/F = (F_response_-F_basal_) / F_basal_) was calculated to represent the effect of shear stress or drugs on calcium signaling.

### RNA interference

The integrin β_3_ siRNA (#sc-63292) and control siRNA (#sc-37007) were purchased from Santa Cruz (Shanghai, China) and the GB1 siRNAs were synthesized from GenePharma (Suzhou, China) and the sequences are: GCGGUUUCCAACGUUCUUUTT (Forward), AAAGAACGUUGGAAACCGCTT (Reverse); GCUACAAGAAGAUCGGCUATT (Forward), UAGCCGAUCUUCUUGUAGCTT (Reverse); GGGAGAAGCCAGUUCCCAUTT (Forward), AUGGGAACUGGCUUCUCCCTT (Reverse).

To knockdown integrin β_3_ expression, HEK293 cells were transfected with lipofectamine 2000 mixed with integrin β_3_ siRNA, with a lipid/siRNA ratio at 2:1. The cells were then cultured for 24 h and subjected to IP_1_ measurements. In parallel, Western blotting was performed to confirm the knockdown of integrin β_3_.

To knockdown GABA_B_ receptor expression, primary astrocytes were transfected with lipofectamine 2000 mixed with GB1 siRNA. The GFP plasmid was co-transfected as reporter of transfection. The lipid/DNA and siRNA ratio was 3:1. Cells were then cultured for 48 h and subjected to Western blotting, calcium imaging and fluorescent imaging. In parallel, quantitative real-time PCR (qPCR) was performed to confirm the knockdown of GB1. The total RNA was extracted with TRIzol (Life Technologies), and qPCR reactions were performed in a 96-well format using Power SYBR Green (Thermo Fisher Scientific). The following primers were used: GB1 forward: ACAGACCAAATCTACCGGGC, GB1 reverse: GTGCTGTCGTAGTAGCCGAT. Relative mRNA levels were calculated using the ΔΔCT method.

### Co-immunoprecipitation (Co-IP)

For HEK293 cells transfected with GABA_B_ receptor with HA-tagged GB1 and Flag-tagged GB2 (HA-GB1 and Flag-GB2), cells were lysed 24 hours after transfection with immunoprecipitation (IP) cell lysis buffer (20 mM Tris, 150 mM NaCl, 1 mM EDTA, 1 mM EGTA, 1% Triton X-100, with complete proteinase inhibitors, pH adjusted to 7.5) on ice for 30 min. For HEK293 cells transfected with GABA_B_ receptor with HA and Snap-tagged GB1, and Flag-tagged GB2 (HA-Snap-GB1 and Flag-GB2), cells were first labeled with 100 nM Snap-649 24 hours after transfection in culture medium at 37 °C for 2 h, then lysed with IP lysis buffer. Lysate were centrifuged at 11,000 g for 10 min at 4 °C. The supernatant was incubated with anti-HA or anti-integrin β_3_ antibody overnight at 4 °C on a rotating wheel and then incubated with Protein G Beads (60 µL of a 50% bead slurry, Millipore) for 4-6 h at 4 °C. The pellet was washed four times with 600 µL of IP cell lysis buffer and resuspended with 2 × SDS sample buffer. Samples were boiled at 95 4 °C for 10 min and subjected to Western blotting analysis.

### Western blotting

Equal amounts of protein were separated by SDS-polyacrylamide gel electrophoresis (PAGE) on 10 to 12% gels. Proteins were transferred to nitrocellulose membranes and washed in blocking buffer (10% BSA in Tris-buffered saline and 0.1% Tween 20) for 2 hours at room temperature. The blots were incubated with primary antibodies overnight at 4°C and then incubated with HRP-linked or fluorescent dye-linked secondary antibodies for 2 hours at room temperature. Immunoblots were detected using Pierce^TM^ enhanced chemiluminescence reagents and visualized by X-ray film, or the membranes were imaged on an Odyssey CLx imager (LI-COR Bioscience, Lincoln, NE, USA) at 700 nm and 800 nm. The density of immunoreactive bands was measured by Image J software and normalized as the fold of the basal control.

### Bioluminescence resonance energy transfer (BRET) titration experiment

BRET titration experiments were performed to measure the direct interaction between proteins as previous reported^57^. Increasing amounts of Venus-tagged receptors were co-expressed with constant amounts of Rluc-tagged receptors in HEK293 cells and cultured for 24 h. Then, after one time wash with PBS, the substrate coelenterazine h (5 μM) was added and BRET signals (emission light at 480 nm and 530 nm, respectively) were detected by Mithras LB940 (Berthold Technologies GmbH & Co., KG). The BRET signals were plotted against the relative expression levels of each tagged receptor. netBRET ratio = [YFP emission at 530 nm/Rluc emission 480 nm] (where Rluc-tagged receptor is cotransfected with Venus-tagged receptor) – [YFP emission at 530 nm/Rluc emission 480 nm] (where Rluc-tagged receptor is transfected alone), in the same experiment. The results were analyzed by nonlinear regression assuming a model with one-site binding (GraphPad Prism, version 8, GraphPad software) on a pooled dataset from three independent experiments.

### Fluorescent imaging

HEK293 cells and astrocytes were fixed with 4% formaldehyde and blocked with 2% BSA and 0.1% Triton X-100 in PBS. HEK293 cells were stained with 4′6-diamidino-2-phenylindole (DAPI) for 30 min. Then cells were washed with PBS and mounted with FluorSave reagent (AR1109, Boster Biological Technology Co. Ltd., Wuhan, China) for fluorescent imaging. Astrocytes were incubated with a primary mouse monoclonal GFAP antibody (1:100) at 4°C overnight. After washing three times with PBS, cells were incubated with a secondary anti-mouse-Cy3 antibody at room temperature for 2 hours. After another washing of three times with PBS, the cells were incubated with DAPI at room temperature for 30 min. Then cells were washed with PBS and mounted with Fluorsave reagent.

Fluorescent images of labeled cells were acquired using an Olympus FV3000 Laser Scanning Confocal Microscope (60 × objective, Olympus, Tokyo, Japan) equipped with appropriate fluorescence and filters (DAPI: 405/449 nm; YFP: 488/519 nm; RFP: 543/620 nm; Cy3: 543/620 nm; DAPI: 405/449 nm; GFP, 488/519 nm). The images were digitized and saved in TIFF format. respectively, more than five microscopic fields were randomly chosen for analysis. Images were processed and fluorescence (yellow in merge panels) was quantified with Image J plugin JACoP using Manders’ co-localization coefficients M1&M2. The settings of the confocal microscope were kept the same for all images when fluorescence intensity was compared. The intensity and area of GFAP were quantified using ImageJ software.

### Intracellular calcium release measurements

The intracellular calcium release measurements were performed as previously described^99^. HEK293 cells were transfected with plasmids encoding the indicated GABA_B_ receptor and a chimeric protein G_qi9_ for 24 h. The cells were then preincubated for 1 h with Ca^2+^-sensitive Fluo-4 (Life Technologies). The fluorescence signals (excitation at 485 nm and emission at 525 nm) were measured for 60 s (Flexstation 3, Molecular Devices) and recorded using a Flexstation 3 microplate reader (Molecular Devices, Sunnyvale, CA, USA). The agonist was added after the first 20 s. The Ca^2+^ response is expressed as an agonist stimulated increase in fluorescence.

### Cross-linking and fluorescent-labeled blot experiments

The Cross-linking experiments were performed as previously described^35^. HEK293 cells were transfected with HA-Snap-GB1^6.56^ and Flag-Halo-GB2^6.56^ by lipofectamine 2000 and plated in 12-well plates for 24 h. Then cells were labeled with 100 nM Snap-649 non cell permeant in culture medium at 37°C for 2 h. Cells were treated in different mechanical force condition with or without GABA (100 μM) treatment. Afterwards, cross-link buffer (1.5 mM Cu(II)-(o-phenanthroline), 1 mM CaCl_2_, 5 mM MgSO_4_, 16.7 mM Tris HCl, pH 8.0 and 100 mM NaCl) was added at 37°C for 30 min. After incubation with 10 mM N-ethylmaleimide at 4°C for 15 min to stop the cross-linking reaction, cells were lysed with lysis buffer (containing 50 mM Tris pH 7.4, 150 mM NaCl, 1% NP-40 and 0.5% sodium deoxycholate) at 4°C for 1 h. After centrifugation at 12,000 × g for 30 min at 4°C, supernatants were mixed with loading buffer at 37°C for 10 min. In reducing conditions, samples were treated without DTT in loading buffer for 10 min before loading the samples. Equal amounts of proteins were resolved by 8% SDS-PAGE. Proteins were transferred to nitrocellulose membranes (Millipore). Membrane was imaged on an Odyssey CLx imager (LI-COR Bioscience, Lincoln, NE, USA) at 800 nm.

### Inter subunit FRET sensor measurement

The pRK5 plasmids encoding GB1b subunit with the extracellular ACP-tag inserted in the lobe1^67^ and N-terminal Snap-tag GB2 subunit^100^ have been described previously. HEK293 cells were co-transfected to express GB1b subunit with the extracellular ACP-tag inserted in the lobe1 and N-terminal Snap-tag GB2 subunit using Lipofectamine 2000 in a 100-mm cell culture dish following the manufacturer’s instructions. Twenty-four hours after transfection, cells were plated in fibronectin (FN)-coated or polyornithine (PLO)-coated white 96-well plates (Greiner Bio-One) at 10^5^ cells per well and cultured 24h at 37 °C and followed by 12 h at 30°C with 5% CO_2_.

To covalently label CoA-Lumi4®-Tb and Snap-Green on GABA_B_ receptors, combined labelling of Snap– and ACP-tag were performed following the protocol described previously with a small modification^101^. Briefly, transfected cells were first incubated with 300 nM Snap-Green in DMEM (1X) + GlutaMaxTM-I medium for 1 h at 37°C, then washed once and incubated with 10 mM MgCl2, 1 mM DTT, 2 μM CoA-Lumi4®-Tb and 1 µM Sfp synthase in Tag-Lite® buffer for 1.5 h at 37°C.

Next, cells were washed three times in Tag-Lite® buffer, and incubated with 60 μL Tag-Lite® buffer. The trFRET measurements were performed in Greiner white 96-well plates, using a PHERAstar FS microplate reader with the following setup; after excitation with a laser at 337 nm (40 flashes per well), the fluorescence was collected at 520 nm. The acceptor ratio was calculated using the sensitized acceptor signal integrated over the time window [50 µs-100 µs], divided by the sensitized acceptor signal integrated over the time window [900 µs-1150 µs].

### Molecular modelling and sequence alignment

The molecular model of GB1 VFT is generated with PyMOL software (Palo Alto, CA, USA) based on the cryo-EM structure of the GABA_B_ receptor (PDB: 7EB2). A model of GB1 VFT to show the residues in LB2 substituted by N-glycan was retrieved from GPCRdb. The protein sequences of the GABA_B1a_ subunit in various species were collected in UniProt (http://www.uniprot.org). The multiple sequence alignment was generated using Clustal Omega (https://www.ebi.ac.uk/Tools/msa/clustalo/) and the graphic was prepared on the ESPript 3.0 server (http://espript.ibcp.fr/ESPript/cgi-bin/ESPript.cgi).

### Statistical analysis

Data are means ± s.e.m. from at least three independent experiments and statistical analyses were performed in GraphPad Prism 8 software. P-values were determined using the unpaired t test (two-tailed), paired t test (two-tailed) or ordinary one-way ANOVA with Tukey’s multiple comparisons test (\*\*\*\**P* < 0.0001, \*\*\**P* < 0.001, \*\**P* < 0.01, \**P* < 0.05 were considered statistically significant). *P* > 0.05 was considered statistically not significant (ns). For dose-response experiments, data are normalized and analyzed using nonlinear curve fitting for the log (agonist) vs. response (three parameters) curves.

## Acknowledgments

We thank the Research Core Facilities of Life Science (HUST, Wuhan) for their assistance in functional measurements. This work was supported by grants from National Key R&D Program of China (grant number 2021ZD0203302 and 2022YFE0116600 to J. L.), the National Natural Science Foundation of China (NSFC) (grant number 32330049, 82320108021 and 32421003 to J. L., 32271198 to C. X.), the Major Project of Guangzhou National Laboratory (grant number GZNL2023A03007 to J.L.) and the Key Technologies R&D Program of Guangdong Province (grant number 2010A080813001 to J. L.).

## Author contributions

J.L., X.Z.S.X. and C.X. conceived and supervised the whole project. Y.Z., and Y.H. performed traction experiments, shear stress experiments, IP_1_ measurements experiments, Co-IP experiments and GB1/integrin interface identification. Y.H. performed fluorescent imaging, BRET experiments and the calcium imaging in astrocytes. L.L., performed the calcium imaging and intracellular calcium release measurements in HEK293 cells, molecular modelling, and sequence alignment. Y.H., Y.Z. and L.L. performed cross-linking experiments, and. F.H. helped to set up the single-cell calcium imaging system. F.Y. helped with the Western blotting detection qPCR, BRET experiment and calcium imaging in astrocytes and HEK293 cells. F.Z. and M.S. helped with the calcium imaging in HEK293 cells. C.S. helped with the molecular modelling. Y.L. helped with the Co-IP and BRET experiment. J.L., X.Z.S.X. and C.X. wrote the manuscript with inputs from all the authors.

## Data Availability

All the other data generated in this study are provided in the Supplementary information and source data files. The raw data for all Figures and Supplementary Figures are available in Source Data files, accompanying this paper. A reporting summary for this article is available as a Supplementary Information file.

## Supplementary figures

**Supplementary Fig. 1.** | Mechanical forces activate GABA_B_ receptor. **a-b**, Immunoblots of phosphorylated MLC2 and β-actin of HEK293 cells under either suspension or adhesion condition *(a)*, or PDL or FN coating *(b)*. Data are present as mean ± s.e.m. from three biologically independent experiments and analyzed using unpaired t test (two-tailed) to determine significance. \*\**P* < 0.01, \**P* < 0.05. **c,** IP_1_ production in cells transfected with vector and G_qi9_, or GABA_B_ receptor and G_qi9_, treated with or without blebbistatin (50 μM, 30 min) under suspension or adhesion condition. Data are present as mean ± s.e.m. from four biologically independent experiments each performed in triplicates and analyzed using unpaired t test (two-tailed) to determine significance. \*\**P* < 0.01, \**P* < 0.05, not significant (ns) > 0.05. **d,** G_i_ protein activation in HEK293 cells transfected with GABA_B_-ΔG a GABA_B_ receptor mutant that fails to couple to G protein, along with G_i_ protein sensors under suspension, PDL-coating or FN coating condition. Data are present as mean ± s.e.m. from six biologically independent experiments each performed in triplicates, and analyzed using repeat measurements one-way ANOVA with Tukey’s multiple comparisons test to determine significance. not significant (ns) > 0.05. **e,** Analysis of the peak value of Ca^2+^ signal in *Fig. 1f*. Data are present as as mean ± s.e.m. from 85 cells and analyzed using ordinary one-way ANOVA with Tukey’s multiple comparisons test to determine significance. \*\*\*\**P* < 0.0001, not significant (ns) > 0.05. **f,** Real-time recording of intracellular Ca^2+^ release in HEK293 cells transfected with vector and G_qi9_. Shear stress, baclofen (100 μM) was applied in the indicated time point. Data are present as mean ± s.e.m. from 239 cells recorded.

**Supplementary Fig. 2.** | Measurement of GABA concentration. **a**, GABA concentration in HEK293 cells transfected with GABA_B_ receptor and G_qi9_ under PDL-coated, FN-coated, or shear stress condition. **b**, Dose response curve of GABA activated Ca^2+^ release in HEK293 cells transfected with GABA_B_ receptor and G_qi9_. The dotting line represent the response of GABA under the concentration in *(a)*. Data are present as mean ± s.e.m. from three biologically independent experiments in *(a)* and *(b)* each performed in triplicates.

**Supplementary Fig. 3.** | GABA_B_ receptor interacts with integrin β_3_. **a**, Expression of integrin β_3_ and GB1 in Fig. 2a. Bars represent the expression of integrin β_3_ and present as mean ± s.e.m. from four biologically independent experiments and analyzed using ordinary one-way ANOVA with Tukey’s multiple comparisons to determine significance. \*\*\**P* < 0.001, \*\**P* < 0.01. **b**, Analysis of the amount of GB1 Co-IPed using anti-IgG or anti-integrin β_3_ antibody in Fig. 2b. Data are present as mean ± s.e.m. from four biologically independent experiments and analyzed using unpaired t test (two-tailed) to determine significance. \*\*\**P* < 0.001. **c-d,** Co-immunoprecipitation of GABA_B_ receptor and integrin β_3_ in HEK293 cells transfected with GABA_B_ receptor (HA-tagged GB1 and Flag-tagged GB2) using anti-integrin β_3_ antibody *(c)* or anti-HA antibody *(d)*. **e,** Co-immunoprecipitation of GABA_B_ receptor and integrin α_v_ in HEK293 cells transfected with GABA_B_ receptor (HA-tagged GB1 and Flag-tagged GB2) using anti-HA antibody. Bars represent the amount of GB1 Co-IPed by anti-integrin β_3_ antibody *(c)*, or integrin β_3_ Co-IPed by anti-HA antibody *(d, e)*. Data are present as mean ± s.e.m. from at least three biologically independent experiments (*c*: n = 4; *d*, n = 4; *e*: n = 3) and analyzed using unpaired t test (two-tailed) to determine significance. **f,** Co-localization of GABA_B_ receptor (YFP fused-GB1 and GB2) and integrin β_3_ fused with RFP in transfected HEK293 cells. (Scar bar: 10 μm, 89.829 ± 1.432% overlapped, n > 20). **g,** Analysis of the amount of integrin β_3_ Co-IPed by anti-HA antibody with or without the following drug treatment: Blebbistatin (50 μM, 30 min), RGDS (10 μM, 12 h) or CGP54626 (50 μM, 30 min) in Fig. 2d-f. Data are present as mean ± s.e.m. from at least three biologically independent experiments (n = 3; 3; 4 for Blebbistatin, RGDS and CGP54626 respectively) and analyzed using unpaired t test (two-tailed) to determine significance. \*\*\*\**P* < 0.0001, \*\*\**P* < 0.001, \*\**P* < 0.01. **h,** Co-immunoprecipitation of GABA_B_ receptor and integrin β_3_ in HEK293 cells transfected with GABA_B_ receptor (HA-tagged GB1 and Flag-tagged GB2) using anti-HA antibody with or without GABA (100 μM, 10 min) treatment. Bars represent the analysis of the amount of integrin β_3_ Co-IPed by anti-HA antibody with or without the GABA treatment. Data are present as mean ± s.e.m. from at four biologically independent experiments and analyzed using unpaired t test (two-tailed) to determine significance. \*\*\**P* < 0.001. **i,** Interaction of GABA_B_ receptor and integrin β_3_ interaction between GB1 and integrin β_1_ detected using BRET titration assay. Data were analyzed by nonlinear regression on a pooled data set from at least three biologically independent experiments each performed in triplicates, fitting with 1-site binding model in GraphPad Prism 8. **j,** IP_1_ production in cells transfected with vector or GABA_B_ receptor along with G_qi9_ chimera under PDL, FN or collagen coating. Data are present as mean ± s.e.m. from three biologically independent experiments each performed in triplicates and analyzed using unpaired t test (two-tailed) to determine significance. \*\**P* < 0.01, \**P* < 0.05. **k,** Co-immunoprecipitation of GABA_B_ receptor and integrin β_3_ in HEK293 cells transfected with GABA_B_ receptor (HA-tagged GB1 and Flag-tagged GB2) or GABA_B_-ΔB mutant (HA-tagged GB1-S246A/E465A and Flag-tagged GB2) using anti-HA antibody under suspension and adhesion condition.

**Supplementary Fig. 4.** | Mapping key residues involved in GABA_B_ receptor and integrin interaction. **a**, Analysis of Co-IPed of integrin β_3_ by anti-HA antibody targeting to GABA_B_-ΔVFT or GABA_B_ in Fig. 4c. Data are present as mean ± s.e.m. from five biologically independent experiments and are analyzed using unpaired t test (two-tailed) to determine significance. \*\*\*\**P* < 0.0001. **b,** Co-immunoprecipitation of GABA_B_-M3 and GABA_B_-M7 with integrin β_3_ in HEK293 cells. The amount of integrin β_3_ Co-IPed by IgG or HA antibody are present as mean ± s.e.m. from five biologically independent experiments and analyzed using ordinary one-way ANOVA with Tukey’s multiple comparisons to determine significance. \*\*\*\**P* < 0.0001, not significant (ns) > 0.05. **c**, Sequence alignment of potential regions involved in GABA_B_ receptor and integrin β_3_ interaction in indicated rat, mouse, human, monkey, bovine, drosophila and C.elegans GB1 subunit with Clustal Omega and displayed by ESPript 3 (http://espript.ibcp.fr/ESPript/cgi-bin/ESPript.cgi). **d,** Co– immunoprecipitation analysis of interaction between GABA_B_-5A and integrin β_3_ in HEK293 cells. The amount of integrin β_3_ Co-IPed by IgG or HA antibody are present as mean ± s.e.m. from four biologically independent experiments and analyzed using unpaired t test (two-tailed) to determine significance. \*\**P* < 0.01.

**Supplementary 5.** | Baclofen activated GABA_B_ receptor to induced astrocyte remodeling. **a**, Representative immunofluorescent staining of GFAP (red) and DAPI (cyan) in astrocytes treated without or with baclofen (100 μM, 14-16h). Scale bar: 10 μm. **b-c,** Analysis of the cell size and GFAP expression in *(a)*. Cell basis (ROI) are presented in the bars as mean ± s.e.m. from indicated number of cells (control: n = 14, baclofen: n = 16) measured in three biologically independent experiments and analyzed using unpaired t test (two-tailed) to determine significance. \*\**P* < 0.01.

**Supplementary Fig. 6.** | GABA_B_ receptor-mediated intracellular calcium release in astrocytes. **a-b**, Real time recording of intracellular Ca^2+^ release in astrocytes transfected with control siRNA *(a)* or GABA_B_ siRNA *(b)* under baclofen (100 μM) and ATP (100 μM) stimulation. Data are representative from three biologically independent experiments. **c,** Percentage of different populations of astrocytes in which baclofen activated calcium response (sensitive) or unable to activate calcium response (insensitive). Data are analyzed with χ2 test to determine significance. \*\*\*\**P* < 0.0001. **d,** Analysis of the peak value of Ca^2+^ signal in Fig. 8a and 8c. Data are present as as mean ± s.e.m. from 15 cells with control siRNA and 45 cells with GABA_B_ siRNA, and analyzed using ordinary one-way ANOVA with Tukey’s multiple comparisons test to determine significance. \*\*\*\**P* < 0.0001, \*\*\**P* < 0.001, \**P* < 0.05.

