## Supplementary figures and images for "GABA-independent activation of GABAB receptor by mechanical forces"

### Supplemental figure 1

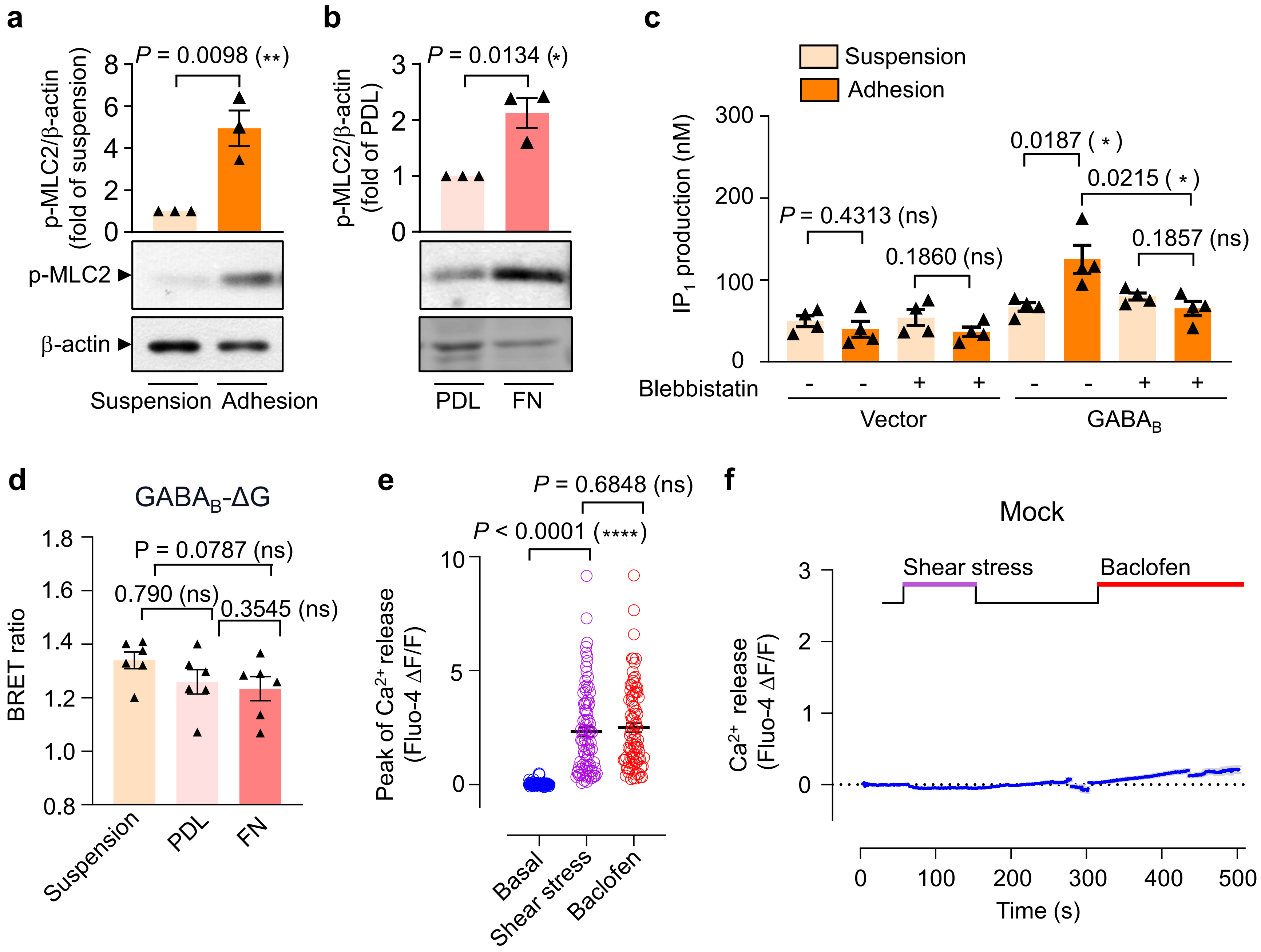

### Supplemental figure 3

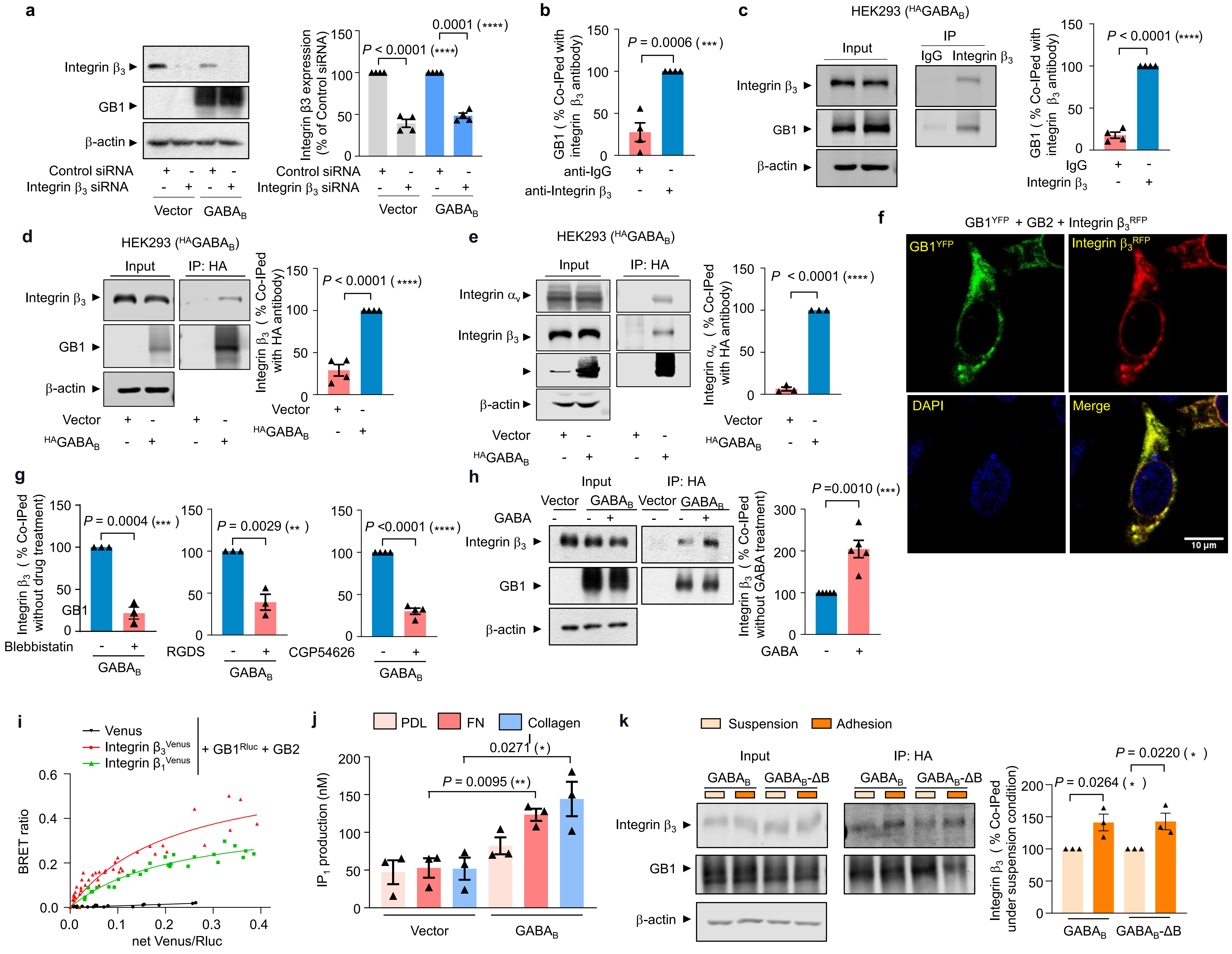

### Supplemental figure 4

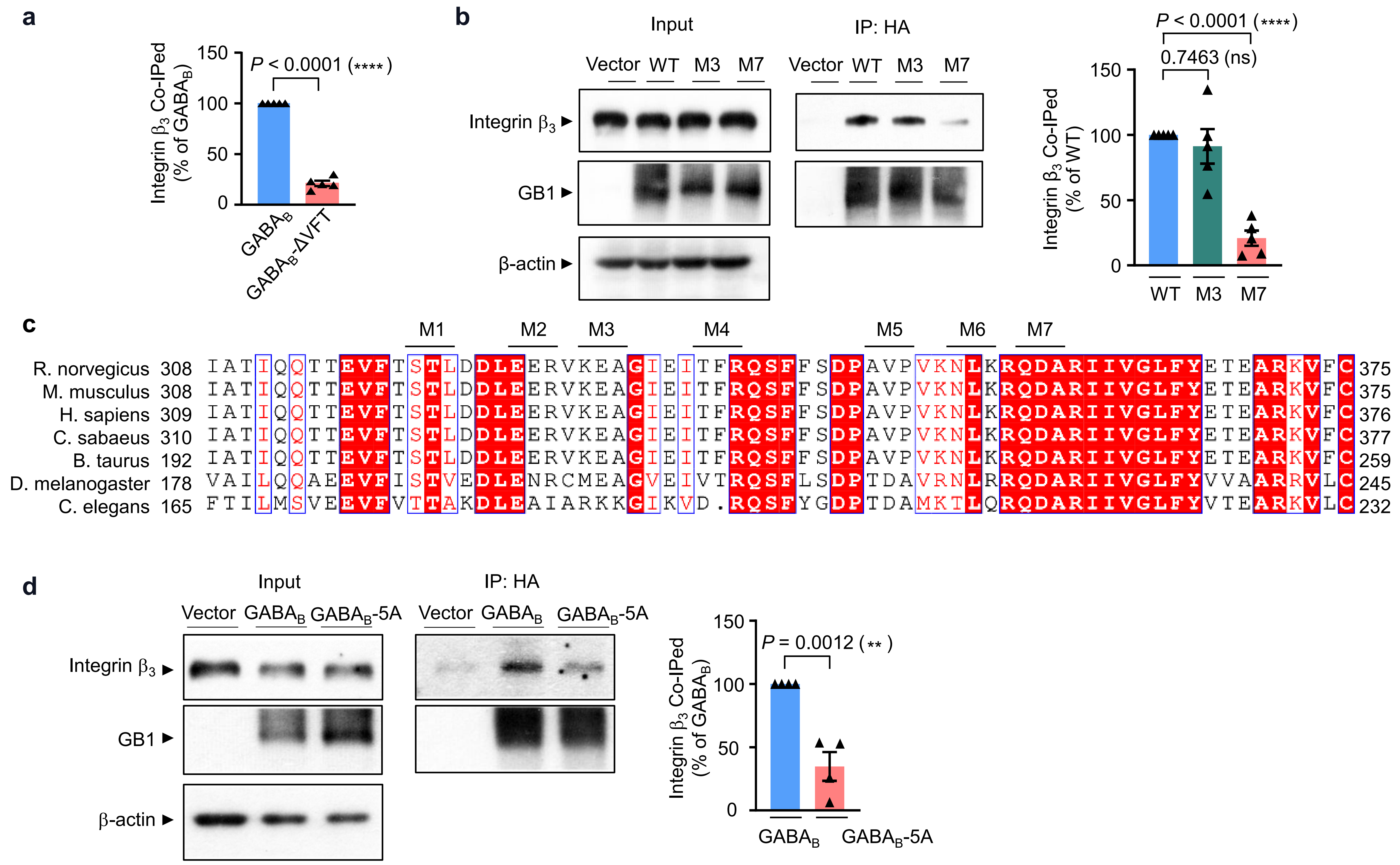

### Supplemental figure 5

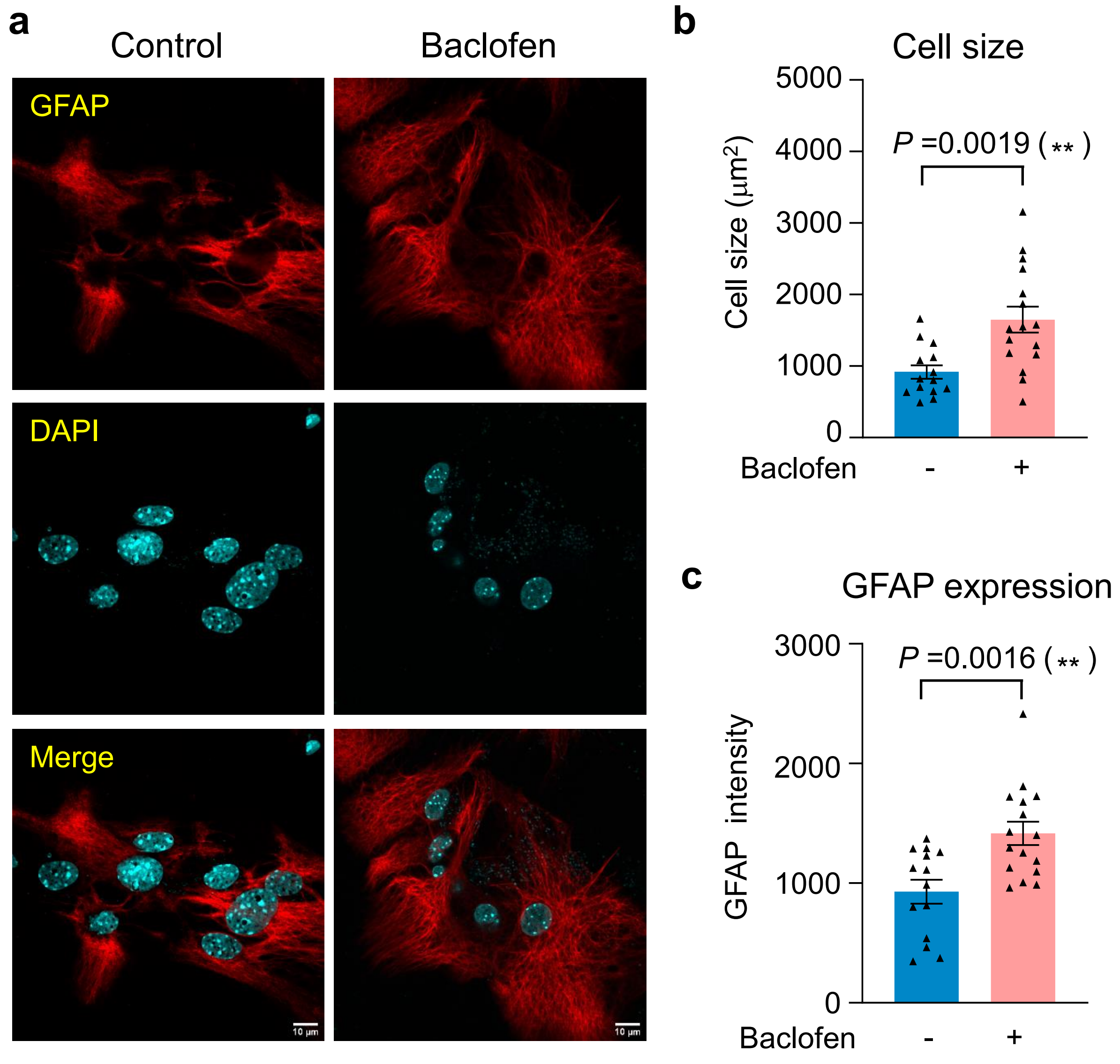

### Supplemental figure 6

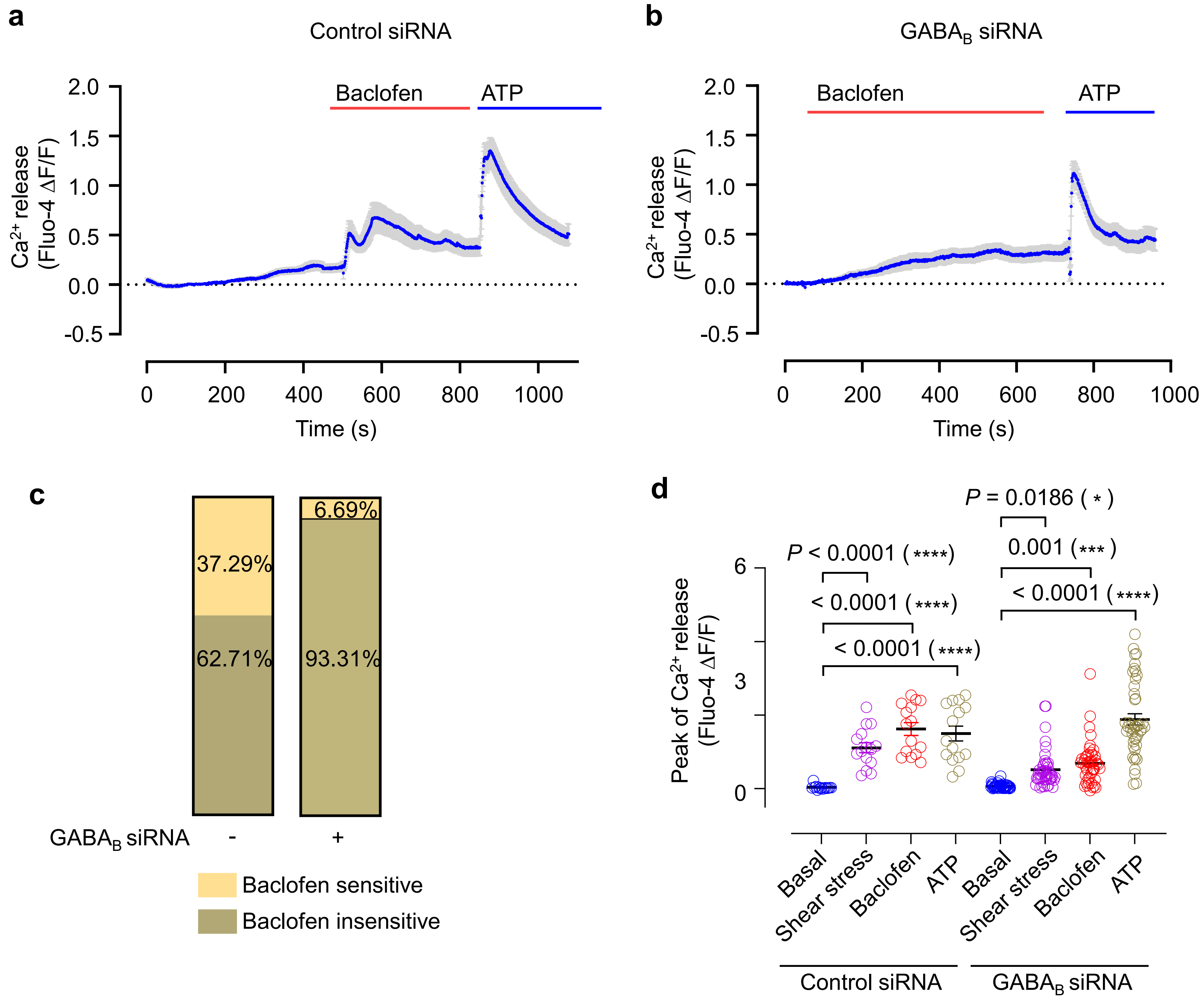
