## Supplemental figure 2 for "GABA-independent activation of GABAB receptor by mechanical forces"

### Figure S2

**a**

HEK293 cells transfected with $GABA_B$ receptor

| | GABA concentration (pM) |
|---|---|
| PDL coated | 20.429 ± 4.297 |
| FN coated | 20.321 ± 5.905 |
| Before shear stress | 18.562 ± 3.250 |
| After shear stress | 16.547 ± 2.246 |

**b**

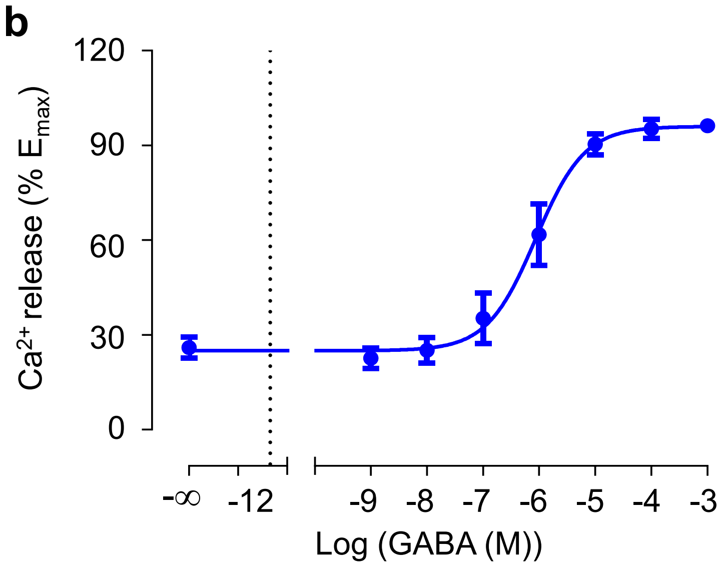
